# Molecular dynamics of Brodmann Area 22 in development and autism

**DOI:** 10.64898/2026.03.31.715694

**Authors:** Varun Suresh, Emilie M. Wigdor, Yuhan Hao, Rachel Leonard, Joseph Asfouri, Michael Griffiths, Clements Evans, Guohua Yuan, Narjes Rohani, Jakob Weiss, Chimmi Dema, Tanzila Mukthar, Frederik H. Lassen, Nicole Schafer, Shan Dong, Duncan S. Palmer, Edward F. Chang, Stephan J. Sanders, Tomasz J. Nowakowski

## Abstract

Challenges in verbal communication are a prominent feature of autism. However, gene regulatory programs in speech-related cortical regions remain poorly characterized. In parallel, it remains unclear whether the heterogeneous genetic factors underlying autism converge on shared neurobiological mechanisms. To address these gaps, we generated paired transcriptomic and epigenomic data from post-mortem human brain tissue across 100 donors. Here, we show that transcriptional differences in the speech-related Brodmann Area 22 in individuals with neurodevelopmental conditions, including autism, are strongest among those with a known genetic diagnosis. A similar but attenuated signature is observed in those without a genetic diagnosis. These transcriptional differences are most pronounced in neurons, with glutamatergic L4/5 intratelencephalic neurons affected across multiple modalities. Finally, multimodal analysis implicates altered *RFX3*-dependent networks as a central hub in autism, particularly among L4/5 intratelencephalic neurons in non-verbal individuals. Together, our study identifies regulatory architecture linking chromatin state, transcriptional output, and variation in verbal ability in autism.

---

Autism spectrum disorder (ASD) is characterized by genetic and phenotypic heterogeneity, including impairments in social communication and language abilities^1,2^. Although ∼15% of autistic individuals carry genetic variants in established ASD-associated and neurodevelopmental disorder (NDD)-associated genes^3^, most liability for ASD arises from common genetic variation distributed across the genome^4–6^. These diverse genetic factors appear to converge on a smaller subset of neurobiological processes, including excitatory and inhibitory neurons, transcriptional regulation, and neuronal communication; however, specific molecular or cellular pathways remain elusive. Molecular profiling of post-mortem human brain tissue has advanced the understanding of ASD by identifying transcriptional and epigenomic differences in the dorsolateral prefrontal cortex, identifying dysregulated programs involved in synaptic development, neuronal connectivity, and circuit maturation^2,7–9^. Molecular changes in language-relevant cortical regions that may underlie differences in verbal abilities are less frequently studied, even though language-related cortical regions may show a larger effect size of differential gene expression and alternative splicing^10^ compared to the prefrontal cortex. Brodmann area 22 (BA22) encompasses Wernicke’s Area and plays an important role in processing auditory information, including language^11–13^, comprehension, and semantic processing; it is consistently implicated in ASD^14,15^. Here, we sought to identify gene regulatory networks underlying differences in ASD and contributing to deficits in language by applying single-nucleus molecular profiling technologies to human post-mortem brain tissue spanning the majority of postnatal life.

## Results

### Multiomic profiling of BA22 across a lifespan in cases and controls

To generate single-cell multiomic profiles of cells in human BA22 from ASD and control individuals, we obtained post-mortem human brain tissue from 100 donors (Fig. 1a; Supplementary Data S1), including controls (N = 52), individuals without a confirmed genetic diagnosis (ASD; N = 34), and individuals carrying established NDD/ASD-associ ated clinically pathogenic/likely pathogenic variants (NDD/ASD; N = 14) (Fig. 1a; Supplementary Data S1-3). We processed single nuclei isolated from pooled tissue of 10–12 donors for snRNA/ATACseq using 10X Genomics technology (Fig. 1b; Supplementary Data S2) to minimize and control for batch effects. In total, we recovered 498,477 single-nuclei with both gene expression and chromatin accessibility profiles passing baseline quality control metrics with a median of 1,766 detected genes and 10,167 ATAC fragments per nucleus (Supplementary Fig. 16-18). Using commonly occurring genetic variation, we demultiplexed individuals, successfully recovering a median of 4,364 nuclei per donor (Supplementary Fig. 1). We assigned cell identities using a combined strategy that integrated Azimuth-based annotation with marker gene validation against previously published datasets^16,17^ (see Methods, Supplementary Fig. 2d-f; Supplementary Data S4). Concordance between automated predictions and manually curated marker profiles enabled confident annotation of all clusters; thus, we identified a total of 26 neuronal and 12 non-neuronal clusters across all donors (Fig. 1c; Supplementary Fig. 2).

**Fig. 1:**
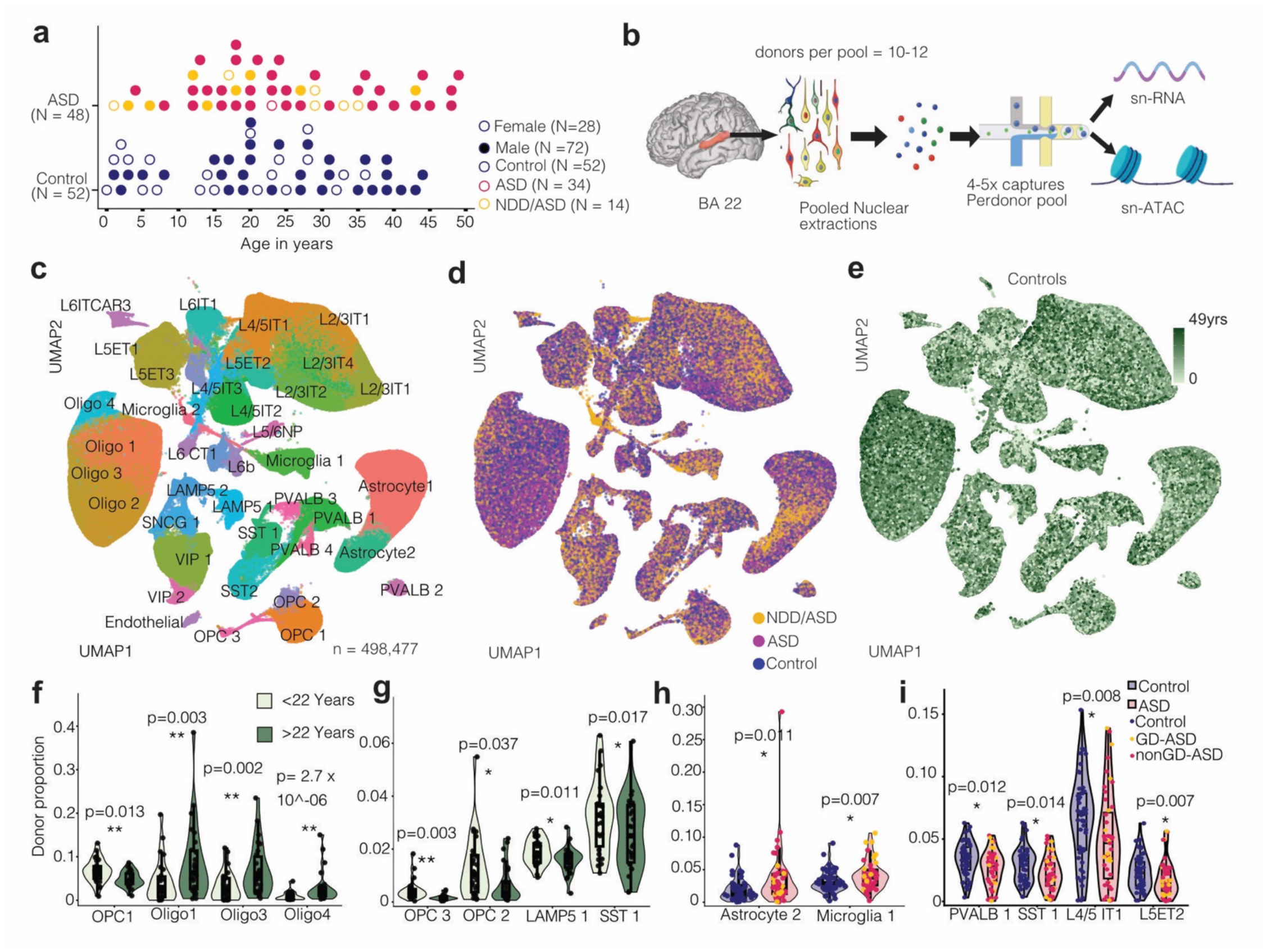
Cohort overview and cellular composition of human BA22 across cases and controls. **a**, Distribution of 100 donors across age and diagnostic categories. Points indicate individual donors, colored by diagnosis (Control, ASD, NDD/ASD) and shaped by sex. **b**, Experimental workflow. 3D rendering of an adult human brain with BA22 (marked in red). Dissected tissue from 10-12 donors was pooled for nuclei extraction, followed by droplet-based multiomic capture (snRNA-seq and snATAC-seq), with 4–5 captures per pool. **c**, UMAP embedding of 498,447 nuclei across all donors, identifying 38 transcriptionally defined cell clusters spanning excitatory neurons, inhibitory neurons, and non-neuronal cell types. **d**, UMAP colored by diagnostic group (Control, ASD, NDD/ASD), demonstrating broad representation of all cell types across groups. **e**, UMAP of control donors colored by age, illustrating representation across the lifespan. **f-g**, Age-stratified cell-type proportion analysis in neurotypical donors (<22 years vs ≥22 years). OPCs decrease with age, whereas mature oligodendrocyte populations increase, consistent with postnatal myelination dynamics. Statistical significance determined using an arcsine-transformed linear modeling approach (Methods). **h-i**, Comparison of cell-type proportions between ASD and control donors. Astrocyte 2 and Microglia 1 clusters show modest increases in ASD, whereas select neuronal subtypes, including L4/5 IT1 and L5 ET2, exhibit moderately reduced proportional abundance. Data points represent proportions calculated in an individual donor. Stars indicate FDR-adjusted p-values; (*) FDR < 0.1, (**) FDR < 0.05.

### Early postnatal cellular composition changes in BA22

To determine if our data could identify changes in cell type proportions by age, we divided cells profiled from control donors younger than 22 years and those older than 22 years and compared cell type proportions using an arcsine-transformed linear modeling framework. We controlled for sequencing depth and total genes detected (Methods) to determine if expected temporal differences in cell type abundance could be detected. We observed a significant (*P* < 0.01, FDR < 0.1) reduction in OPC populations and a reciprocal increase in mature oligodendrocyte clusters in older donors (Supplementary Fig. 3a; Fig. 1e-g), consistent with the expected changes related to delayed myelination. Two interneuron clusters, SST1 and LAMP5-1, showed modest reductions (*P* < 0.05, FDR < 0.1) with age (Fig. 1g, Supplementary Data S5), while no other major cell classes displayed significant proportional shifts (Supplementary Fig. 3c). To further resolve maturation-related changes in interneuron proportions, we performed iterative subclustering of adult interneurons using the hicat algorithm^18^, followed by sequential label transfer to younger age groups. All subclusters were represented across age groups (Supplementary Fig. 4a). Subcluster-level analysis revealed increases in SST_MYO5B, SST_ERBB4, SNCG_PENK, and SST_SPHKAP populations with age (*P* < 0.05, FDR < 0.1), alongside decrease s in VIP_SMOC1 and SST_SYT9 (P < 0.05, FDR < 0.1); Supplementary Fig. 3c–i; Supplementary Data S5), indicating interneuron subtypes undergo transcriptomic refinement during postnatal maturation. Finally, we compared proportions of excitatory neurons, inhibitory neurons, astrocytes, oligodendrocytes, and OPCs across all cases and controls. We found an increased proportion of astrocytes and microglia in cases (Supplementary Fig. 3b), as previously reported^19^ (*P* < 0.05, FDR < 0.1), and these changes were accompanied by reductions in select neuronal populations, including L4/5 IT1, L5 ET2, PVALB 1, and SST 1 (*P* < 0.05, FDR < 0.1; Fig. 1h, i; Supplementary Data S5).

### Early postnatal transcriptional dynamics in BA22

To identify the dynamics of BA22 gene expression, we employed variance partitioning using dreamlet to identify 27,585 unique age-associated genes across 25 clusters (Methods), with the largest numbers observed in excitatory and inhibitory neurons (Supplementary Fig. 5a). Analysis of gene expression trajectories revealed rapid changes in gene expression in the first few years of life (Supplementary Fig. 5b,c), consistent with prior reports^20,21^. These profiles could be grouped into six co-expression modules (Supplementary Fig. 5d). Several genes exhibited conserved trajectories across neuronal and glial populations. For example, *PLPPR4, LUZP2,* and *SPHKAP* declined with age, whereas *STAT4* and *ZBTB16* increased (Supplementary Fig. 5f–g). Language-associated genes showed distinct temporal signatures: *FOXP1, FOXP2,* and *ROBO1* followed Early-fall or Fall patterns, whereas *CNTNAP2, DCDC2,* and *ROBO2* exhibited Early-rise or Late-peak dynamics (Supplementary Fig. 5h, i), indicating structured transcriptional transitions during early childhood. The overall rates of transcriptional changes declined and stabilized across cell types by early childhood (Supplementary Fig. 5 b-d). Together, our analyses demonstrate that previously described dynamic remodeling of cell type specific gene expression profiles during early postnatal development of the prefrontal cortex are observed in BA22.

### Transcriptional dysregulation across neuronal populations in NDD/ASD

To characterize transcriptional differences in cases, we identified differentially expressed genes (DEGs) across 37 neuronal and non-neuronal cell types (Methods) (Supplementary Data S6). We first compared all cases (N=48; “All”) to controls (N=52). We identified 1,776 unique DEGs (FDR < 0.05) in the “All” comparison, 75.5% of which were unique to a single cell type (Fig. 2a). Subsequently, when we restricted the comparison to only NDD/ASD cases, we identified 7,828 unique DEGs, 38.5% of which were unique to one cell type. Excitatory neuron subclasses exhibited the largest number of DEGs, particularly deep-layer IT and ET populations. The extensive number of neuronal DEGs is consistent with prior single-cell NDD/ASD studies^19^. In contrast to previous studies, we did not detect a large number of DEGs within glia^22,23^, possibly reflecting differences in cohort size, statistical power, and batch correction. Consistent with prior work, we found that DEGs had significantly increased odds of being ASD-associated genes (All: OR = 2.07, *P* = 2.22 x 10^-10^ and NDD/ASD: OR = 1.65, *P* = 7.22 x 10^10^) (Methods) and NDD-associated genes (All: OR = 2.30, *P* = 5.71 x 10^-5^ and NDD/ASD: OR = 1.55, *P* = 3.72 x 10^-3^) (Fig. 2c)^3^. Downregulated genes were enriched for mitochondrial, oxidative phosphorylation, ATP metabolic, and ribosomal processes (*P* < 1 × 10^-20^), whereas upregulated genes were enriched for neuronal projection development, dendrite morphogenesis, synapse organization, cell junction assembly, and small GTPase signaling (*P* < 1 × 10⁻^9^) (Fig. 2d–e; Supplementary Data S7).

**Fig 2.**
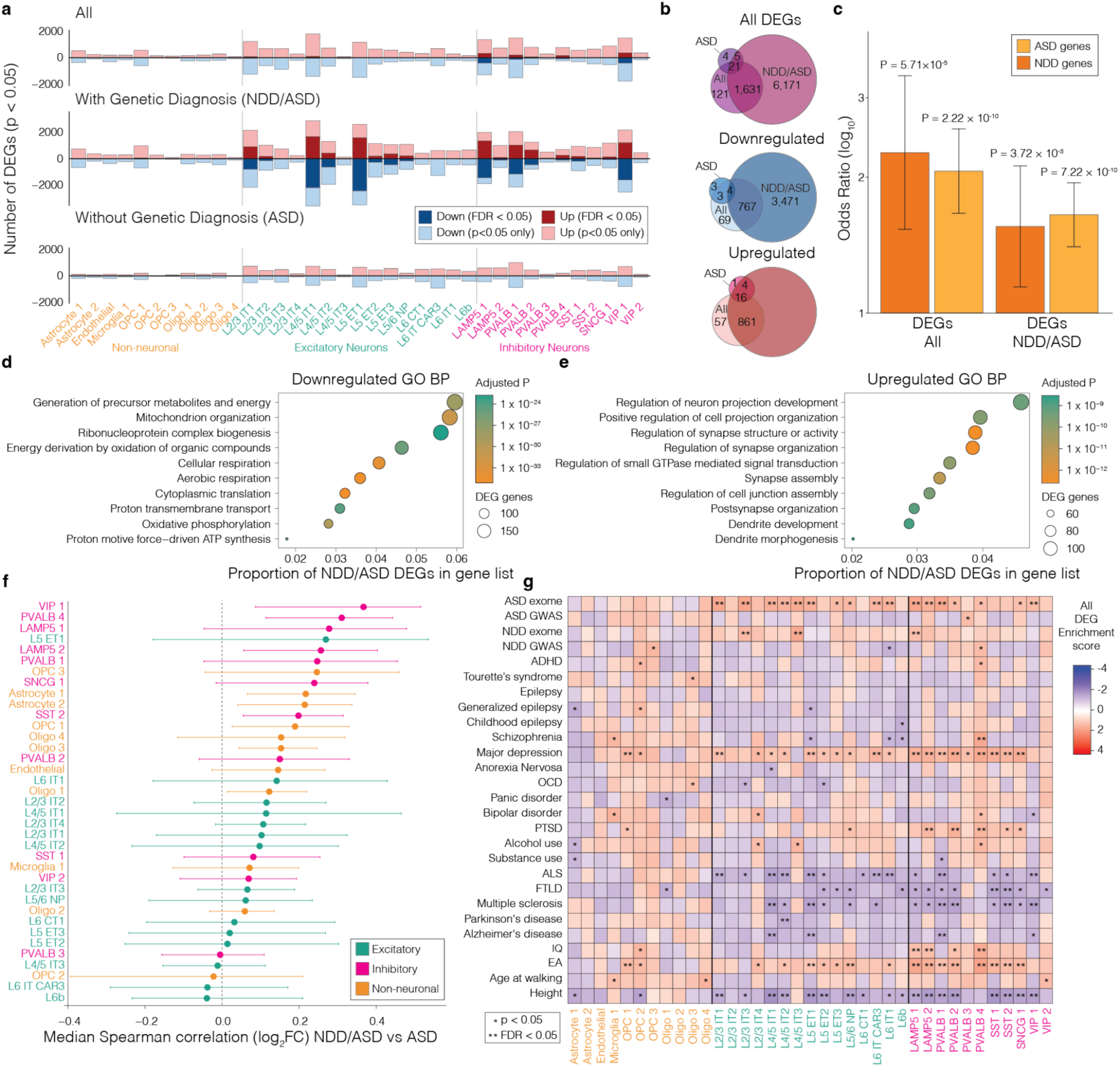
Differential expression and gene set enrichment in autism with and without a genetic diagnosis. **a**, Number of differentially expressed genes (DEGs) per cell type. Cell types are grouped as non-neuronal, excitatory neurons, and inhibitory neurons. Dark red/blue bars indicate FDR-significant (FDR < 0.05) up- and downregulated genes, respectively; light red/blue bars indicate nominally significant genes (*P* < 0.05). Rows correspond to comparisons: All (all cases vs controls), NDD/ASD (cases with genetic diagnosis vs controls), and ASD (cases without diagnosis vs controls). DEGs were identified using precision-weighted linear mixed models (dreamlet; Methods). **b**, Overlap of significant DEGs across comparisons (All, NDD/ASD, ASD) for all DEGs combined, and separately for upregulated and downregulated genes. **c**, Enrichment of DEGs (All and NDD/ASD) for autism risk genes and NDD genes identified via rare variant burden^3^. Odds ratios (log_10_ scale) and 95% confidence intervals are shown; p values from Fisher’s exact test. **d–e**, Gene Ontology (GO) biological process enrichment among downregulated **d**, and upregulated **e**, DEGs from the NDD/ASD comparison. P-values are Benjamini–Hochberg corrected from hypergeometric tests using all expressed genes as background. **f**, Concordance of differential expression between NDD/ASD and ASD comparisons. Control donors were randomly split 50/50; log_2_FCs were estimated independently and correlated across all expressed genes. Procedure repeated 1,000 times; points show median Spearman’s ρ and 95% confidence intervals. **g**, Gene-set enrichment analysis (fGSEA) by cell type for DEGs from all cases vs controls. Gene sets include the top 500 GWAS genes (Methods) and rare variant gene lists for autism and NDD^3^ ( * *P* < 0.05; ** FDR < 0.05.)

### Transcriptomic signatures dysregulated in NDD/ASD correlate with ASD only

Next, we repeated the analysis across donors without a confirmed genetic diagnosis (ASD cases) and identified only 33 DEGs, despite having a larger sample size than the NDD/ASD group. To test whether the transcriptomic signatures identified in NDD/ASD cases extended to individuals without a genetic diagnosis, we calculated the correlation of the log_2_ fold-changes (log_2_FC) of genes from the NDD/ASD and ASD DGE analysis. To obtain an estimate of concordance in differential expression across subgroups, we calculated correlations using all expressed genes rather than restricting to FDR-significant hits, which avoids the winner’s curse inflation associated with thresholding. In addition, because shared controls can induce artificial similarity between contrasts, we repeatedly (1,000 iterations) split the control donors into independent subsets when computing log_2_FCs (Methods; Fig. 2f). We found that inhibitory neurons showed the highest median correlations in effect sizes (Spearman’s ρ 0.38 - 0.26), whereas excitatory neurons did not (Fig. 2f). This analysis suggests that ASD donors (without a genetic diagnosis) show modest concordant gene expression changes with NDD/ASD individuals, but with smaller effect size, particularly among cortical GABAeric neurons.

### DEGs enriched for neuropsychiatric and neurodevelopmental traits

As many neuropsychiatric and neurodevelopmental traits are highly genetically correlated^24–28^, we assessed whether genes associated with these conditions via genomic studies, including genome-wide association studies (GWAS)^3,27,29–51^ and exome sequencing studies (ASD and NDD)^3^ were enriched toward the top or bottom of gene lists ranked by differential expression in each cell type (NDD/ASD). Significant (FDR < 0.05) enrichment was highly cell-type specific and concentrated in neuronal populations (Fig. 2g), including both excitatory projection neurons (L2/3–L6 IT/ET/NP) and inhibitory interneurons (SST, PVALB, VIP) for multiple neuropsychiatric traits (Fig. 2g; Supplementary Data S8). Upregulated DEGs were enriched for ASD genes identified through exome sequencing, whereas enrichment for similarly identified NDD genes was more limited and observed in only three cell types. Upregulated DEGs were also enriched for major depressive disorder^39^, educational attainment (EA)^42^, cognitive performance (IQ)^44^ and post-traumatic stress disorder (PTSD)^40^ (Fig. 2g). In contrast, downregulated DEGs were enriched for neurodegenerative disorders, including ALS, FTLD, multiple sclerosis, Parkinson’s disease, and Alzheimer’s disease, particularly within excitatory neuronal populations. Similar enrichment patterns were observed across all cases and controls (All), as well as in ASD cases only (Fig. 2g; Supplementary Fig. 6). These findings suggest convergence of common neuropsychiatric genetic risk on neuronal transcriptional programs, with distinct directionality relative to neurodegenerative disease-associated loci.

### Motif accessibility differences implicate *RFX3* and immediate early genes in ASD

To investigate ASD-associated changes in chromatin accessibility, we used snATAC data to define differentially accessible regions (DARs) by cell type (Methods) (Supplementary Data S9). Similar to the DEGs from transcriptomic analysis, DARs were almost exclusively identified in NDD/ASD donors (N = 1,101; 98.7% of unique DAR-cell type pairs across comparisons) (Fig. 3a; Supplementary Fig. 7-9), and few DARs passed multiple testing across all cases and controls (N = 7), or in ASD donors alone (N = 7) (Supplementary Fig. 9). More peaks showed reduced accessibility (N = 1,033) than increased accessibility (N = 68) in cases, with excitatory neurons (L2/3 IT1 and 2, L4/5 IT1 and 2) showing the strongest overall effect size (Fig. 3a; Supplementary Fig. 7).

**Fig 3.**
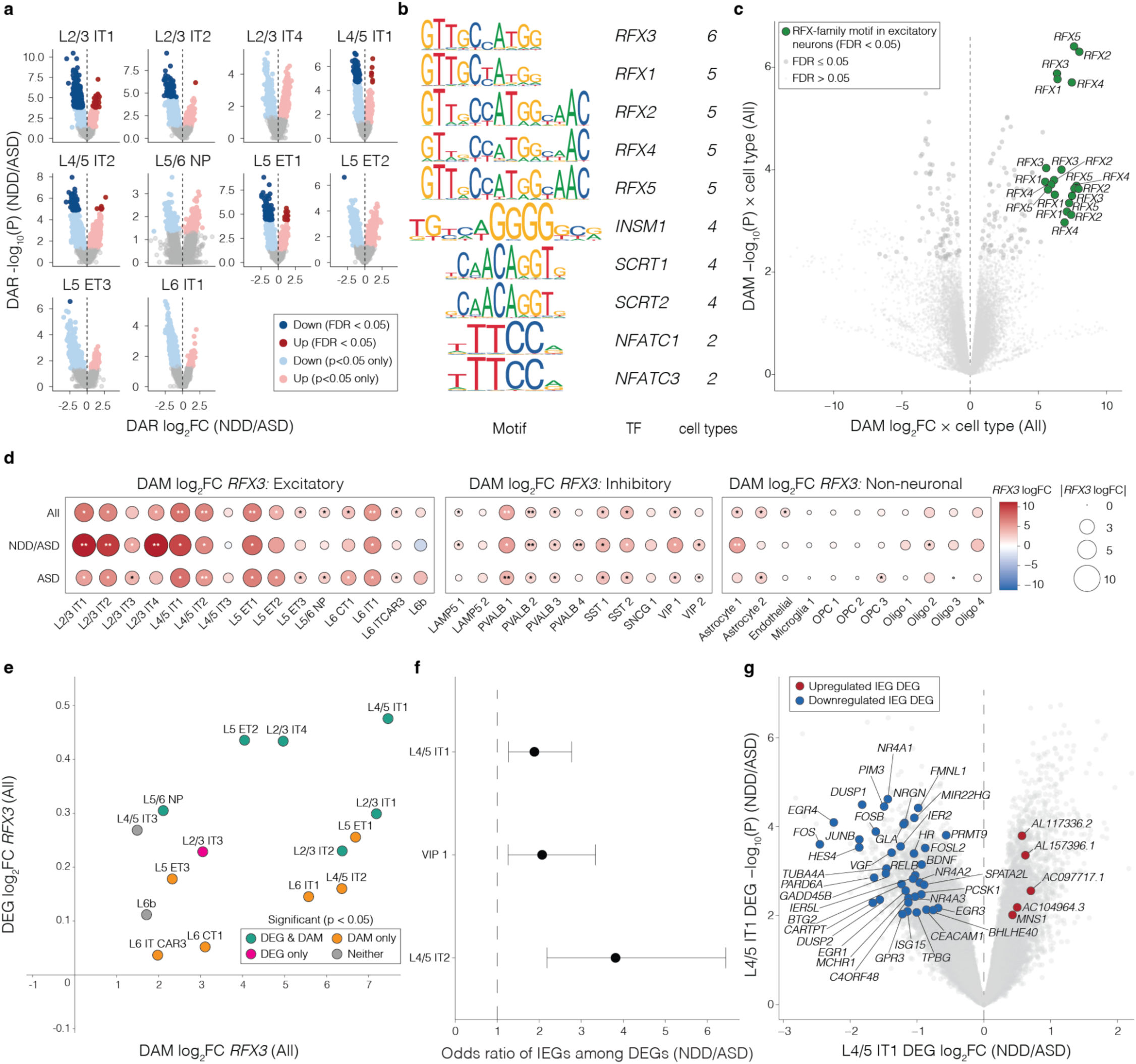
Cell-type-specific chromatin accessibility implicates *RFX3* dysregulation in autism. **a**, Differentially accessible regions (DARs) in NDD/ASD donors. Each point represents an ATAC-seq peak. Dark red and blue indicate FDR-significant (FDR < 0.05) increased and decreased accessibility in cases, respectively; lighter colors indicate nominal significance (*P* < 0.05). **b**, Top 10 transcription factor motifs with recurrent differential accessibility across cell types (All comparison). Motifs and the number of cell types with FDR-significant differential accessibility are shown. **c**, Differentially accessible motifs (DAMs) across cell types. Each point represents a motif-cell type pair; green indicates RFX-family motifs in excitatory neurons. DAMs were identified using linear models implemented in limma (Methods). **d**, *RFX3* differential motif accessibility (log_2_FC) across excitatory, inhibitory, and non-neuronal cell types for All, NDD/ASD, and ASD comparisons. **e**, Relationship between differential *RFX3* motif accessibility and differential *RFX3* expression in excitatory neurons (NDD/ASD). Points colored by nominal significance in accessibility and/or expression. X-axis is the log_2_FC from differential motif accessibility analysis and the y-axis is the log_2_FC for differential gene expression analysis. **f**, Enrichment of immediate early genes (IEGs) among DEGs in significant (FDR < 0.05) cell types (NDD/ASD). Odds ratios and 95% confidence intervals from Fisher’s exact test. **g**, Differential expression of IEGs in L4/5 IT1 neurons (NDD/ASD). Red and blue indicate FDR-significant (FDR < 0.05) increased and decreased expression in cases, respectively.

To identify potential regulatory drivers, we assessed changes in motif accessibility by cell type (Methods). The motif of *RFX3*, an ASD-associated gene from exome sequencing (FDR 3x10^-7^) ^3^, was the most consistent differentially accessible motif (DAM) across cell types in the ‘All’ comparison, showing increased accessibility in cases relative to controls (FDR < 0.05 in 6 cell types; *P* < 0.05 in 13 additional cell types; Supplementary Data S10). In the NDD/ASD comparison, *RFX3* ranked second (tied with *RFX4* and *RFX5*) by recurrence across cell types after *RFX2* (Supplementary Fig. 10; Supplementary Data S10). In the ASD only comparison, *RFX3* was among the most recurrent DAMs, tied for the highest count (2 cell types), although multiple TFs shared this level of recurrence (Fig. 3b–c; Supplementary Fig. 10; Supplementary Data S10). Similar to the DARs, *RFX3* motifs were most strongly enriched among L2/3 IT, L4/5 IT, and L5 ET neurons (Fig. 3d). The magnitude and direction of *RFX3* motif accessibility were consistent across cell types for all three case comparison groups (Fig. 3d). Given the differential accessibility of *RFX3* motifs, we considered whether *RFX3* expression was also altered in excitatory neurons (Fig. 3e). Across all excitatory neuron subtypes, both *RFX3* expression and *RFX3* motif accessibility were increased, with one or both reaching significance thresholds in most subtypes (All comparison; *P* < 0.05). This suggests coordinated regulatory and transcriptional dysregulation (Fig. 3e), with the greatest differences observed in L4/5 IT1. Consistent patterns were observed in the NDD/ASD and ASD comparisons (Supplementary Fig. 11).

*RFX3* has been shown to increase the transcription of activity-dependent immediate-early gene (IEGs) by enhancing CREB binding in neurons^52,53^. Given the increased *RFX3* expression and increased *RFX3* motif accessibility in cases, we investigated whether differential *RFX3* activity correlated with altered expression of IEGs. In the NDD/ASD comparison we found that IEGs were significantly (FDR < 0.05) enriched among DEGs in L4/5 IT1 (OR = 1.88; adjusted *P* = 0.02, Fisher’s exact test) and L4/5 IT 2 (OR = 3.82; adjusted *P* = 1.14 x 10^-4^), as well as in VIP 1 inhibitory neurons (OR = 2.07; adjusted *P* = 0.03) (Fig. 3f). Of note, the differentially expressed IEGs showed predominantly decreased expression (Fig. 3g), raising the possibility that *RFX3* changes are compensatory.

### Cell-type–specific regulatory programs associated with verbal phenotypes in ASD

Given the role of BA22 in human verbal communication, we sought to identify the key cell types and molecular signatures associated with verbal/non-verbal status in the donors. Phenotype data on verbal communication were available for a subset of individuals (four NDD/ASD and seven ASD donors were non-verbal, four NDD/ASD and 26 ASD donors were verbal or unrecorded for any verbal phenotypes, see Methods); non-verbal status was not associated with a genetic diagnosis (*P* = 0.18, two-sided Fisher’s Exact test). To minimize potential confounding effects of sex and age (Fig. 1), we further restricted the analysis to male individuals over two years old (as all non-verbal donors were male, precluding robust modelling of sex as a covariate). First, we identified a nominally significant (*P* < 0.05) increase in the relative proportion of PVALB 1 and L4/5 IT1 neurons in non-verbal compared to verbal cases (Fig. 4a, Supplementary Fig. 12a). These differences were not identified when we compared individuals with versus without intellectual disability (Supplementary Data S11), suggesting that alterations in specific neuronal subtypes are associated with language phenotypes rather than general cognitive impairment.

**Figure 4.**
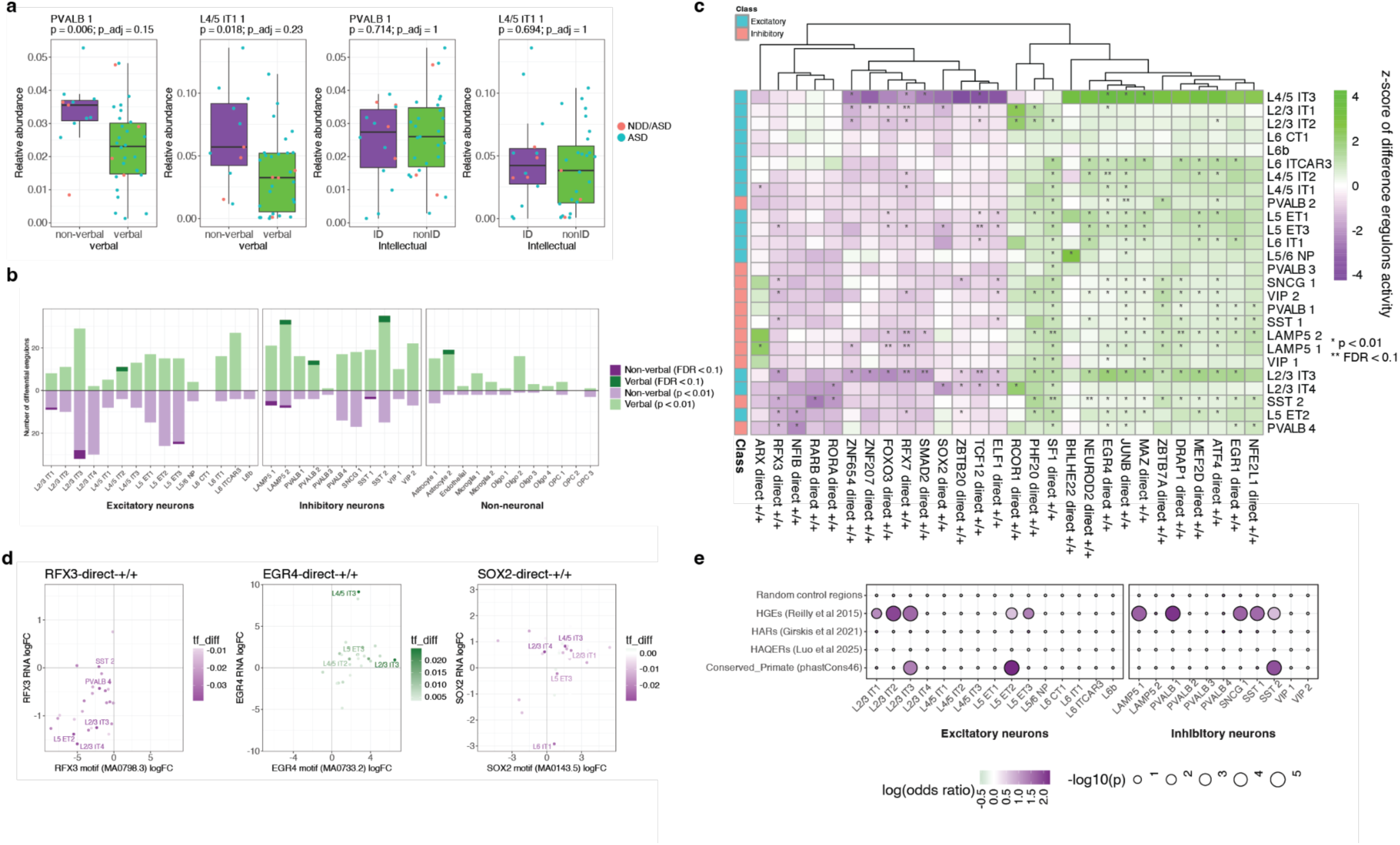
Cell-type specific regulatory programs associated with verbal phenotypes in ASD. **a**, Relative proportions per donor of PVALB 1, L4/5 IT 1 clusters in non-verbal versus verbal ASD cohorts and individuals with versus without intellectual disability. NDD/ASD patients are shown in red dots, and ASD patients are shown in blue dots. P-values and false discovery rate (FDR) adjusted P-values were calculated using two-sided Mann-Whitney U tests. **b**, The number of differential eRegulons calculated by SCENIC+ across excitatory, inhibitory, and non-neuronal lineages. Green and purple bars denote eRegulons with significantly higher activity in verbal and non-verbal ASD, respectively (light shades: *P* < 0.01; dark shades: FDR < 0.1; Mann-Whitney U test). **c**, Hierarchical clustering heatmap of z-scored differences in activity for positive eRegulon drivers across neuronal subtypes. Included eRegulons met the significance threshold of *P* < 0.001 in at least one neuronal subtype. Asterisks indicate statistical significance (*P* < 0.01), and double asterisks indicate FDR-adjusted significance (FDR < 0.1). **d**, Multi-omic coordination of TF expression, motif accessibility, and eRegulon activity. Scatter plots illustrating the coupling between differential TF mRNA expression (y-axis) and matched DNA-binding motif accessibility (x-axis) between verbal and non-verbal ASD patients. Points represent individual cell types and are colored by the magnitude of inferred differential eRegulon activity. Representative drivers reveal distinct regulatory patterns: RFX3 exhibits cell-type-dependent differential activity across L2/3 IT 3, L5 ET 2, and L4/5 IT 1 populations, while EGR4 shows pronounced differential activity specifically within L2/3 IT 3 neurons of verbal individuals. SOX2 is used as a control. **e**, Enrichment of differential eRegulon target peaks within evolutionary and functional genomic annotations, including fetal brain human-gained enhancers (HGEs), human accelerated regions (HARs), and human ancestor quickly evolved regions (HAQERs). Dot size reflects statistical significance (−log_10_ P-value) and color intensity represents the log(odds ratio) derived from Fisher’s exact tests.

To identify transcriptomic and epigenomic features associated with non-verbal status, we performed differential activity analysis of enhancer-driven gene regulatory networks (eRegulon) inferred by SCENIC+. We identified widespread differential eRegulon activity, predominantly in neurons (Fig. 4b, Supplementary Fig. 12b,c). Both cell-type specific and shared differential regulons were observed, including transcription factors such as *EGR4* and *RFX3* (Fig. 4c,d). Notably, *RFX3* exhibited differential regulatory activity not only between ASD and control individuals, but also within the ASD cohort when comparing verbal and non-verbal groups (Supplementary Data S12). This effect was cell-type dependent: L2/3 IT3, L2/3 IT4, and L5 ET2 neurons showed coordinated differences in *RFX3* RNA expression, motif accessibility, and eRegulon activity.

Finally, considering that language is a recently evolved trait, we integrated recently evolved genomic loci, including human accelerated regions (HARs)^54^, human ancestor quickly evolved regions (HAQERs)^55^, fetal brain human-gained enhancers (HGEs)^56^, and primate-conserved regions (Methods)^57^. HGEs and primate-conserved regions have previously been shown to be enriched for genetic heritability associated with functional language networks and vocabulary ability^58^. We tested whether chromatin accessibility peaks that belonged to eRegulons enriched in non-verbal cases (Fig 4 b,c) were over-represented for highly-evolved genomic regions. We observed a significant enrichment of eRegulon-associated peaks within both HGEs and primate-conserved regions (Fig. 4e). Notably, altered open-chromatin peaks in L2/3 IT3, L5 ET2, and SST2 neurons were enriched across both categories, indicating overlap between ASD-associated regulatory changes and recently evolved genomic elements linked to human language-related genomic regions.

## Discussion

Impairments in verbal communication are a common feature of ASD. Due to the uniqueness of sophisticated language to the human species, the neurobiological mechanisms underlying language-related phenotypes are difficult to study in experimental systems. Here, we performed multiomic profiling of human BA22 from post-mortem tissue across the postnatal developmental trajectory in individuals with ASD and controls. We define cell type specific transcriptional changes underlying the early development of BA22, a cortical region involved in auditory processing, including speech information.

Within the first years of life, the cell type composition of BA22 changes, most notably among transcriptomic subtypes of cortical interneurons. In addition, cell types undergo dynamic changes in gene expression programs that include changes in the expression of transcription factors implicated in language through disease studies. Similar early developmental shifts have been observed across multiple cortical regions^20,21^, suggesting that dynamic transcriptional remodeling during early postnatal development may represent a broadly conserved feature of human cortical maturation.

Across molecular layers, the ASD-associated alterations in BA22 we observed fall into three broad categories: transcriptional changes, chromatin accessibility differences, and changes in cell type abundance. The magnitude of molecular disruption is substantially greater in donors carrying established NDD/ASD-associated pathogenic variants. This pattern is consistent with the expectation that high-penetrance variants produce larger transcriptional effects detectable in post-mortem steady-state tissue. A key question is whether transcriptional signals converge on similar pathways between individuals with and without a detected genetic diagnosis. Although few genes reached statistical significance in the ASD comparison, effect sizes were partially concordant with those observed in genetically diagnosed individuals, especially among GABAergic cortical interneurons. These data support a molecular spectrum model in which monogenic or high-penetrance variants produce broad, high-amplitude perturbations, whereas in the absence of such variants, cases engage overlapping regulatory programs with smaller, more heterogeneous effects that will require larger cohorts to resolve fully. Our findings are broadly consistent with prior bulk and single-cell studies reporting neuron-centered dysregulation and perturbation of synaptic and activity-dependent programs in ASD, but as a notable exception, we did not find strong evidence of dysregulated glial transcriptomes^10,19,22^.

Integration of chromatin accessibility, motif analysis, gene expression, and enhancer-driven regulon inference identifies recurrent perturbation of the *RFX* transcription factor family across neuronal populations. *RFX* genes have been implicated in multiple psychiatric disorders and regulate neuronal cell fate and developmental processes through cooperative binding to DNA, often with multiple RFX family members in combination with other transcription factors^59–61^. Among these, *RFX3* stands out. *RFX3* is a transcription factor critical for cortical development and a high-confidence ASD risk gene^3,59,60,62^. Interestingly, *RFX3* has been shown to control immediate early gene expression via the cAMP-response element-binding protein (CREB) pathway in experimental model systems, implicating activity-dependent signaling or altered cofactor interactions in autism^52,53^. IEG programs have long been proposed as central mediators of activity-dependent transcriptional responses in neurodevelopmental disorders^63^, and a key CREB binding protein (*CREBBP*) is also associated with ASD^3^. *RFX3* motif accessibility is increased across multiple excitatory subtypes; its transcript levels are elevated in several of these populations, most prominently among L4/5 IT1 neurons, a set of cortico-cortical projection neurons that integrate local sensory input and distribute information across cortical networks, making them potential components of association circuits involved in language processing.

Our analysis of verbal versus non-verbal individuals within the ASD and NDD/ASD cohorts revealed cell-type specific alterations in gene regulatory networks that were associated with language ability, rather than general cognitive impairment. Differential eRegulon analysis identified transcription factors such as RFX3 with coordinated changes in expression, motif accessibility, and regulatory activity across specific BA22 neuron subtypes, suggesting that disruption of precise gene regulatory programs in defined neuronal populations may contribute to the profound language deficits observed in a subset of individuals with ASD. Strikingly, the regulatory regions associated with these differential eRegulons were enriched within human-gained enhancers and primate-conserved genomic regions previously linked to language-related traits in population-level analyses^58^. This raises the possibility that the interplay between neurodevelopmental dysregulation and human-specific evolutionary innovations at these regulatory elements contributes to the diverse language outcomes observed in ASD.

Several limitations warrant consideration. Early postnatal sampling remains limited and may be particularly informative for resolving when molecular differences first emerge across ASD/NDDs and questions of cause versus consequence. Post-mortem intervals may alter the transcriptome such that our data differ from the living state. The genetically diagnosed subgroup is heterogeneous with respect to variant class and possible downstream mechanisms. In the genetically undiagnosed group, sex imbalance (only a single female donor) precluded inclusion of sex as a covariate, necessitating male-only analyses and thereby reducing statistical power and generalizability. Clinical annotation, including verbal phenotype data, is incomplete for some donors and may introduce bias in analyses comparing verbal and non-verbal cases if missingness is systematic. In addition, DEG and DAR burden estimates are influenced by cell-type specific power, transcriptome complexity, cohort composition, and analytical choices, including model selection and covariate adjustment, which can alter the number of detected features^64^; absolute counts should therefore be interpreted cautiously (Supplementary Fig. 13). Similarly, interpretation of GWAS enrichment should consider differences in statistical power across traits, as current studies vary substantially in sample size; enrichment patterns may therefore shift as larger genetic datasets become available. Finally, we note that a subset of donors had comorbid epilepsy, and detailed medication histories were not uniformly available. Seizure activity or anti-epileptic treatment could plausibly influence activity-dependent transcriptional programs, including IEG expression; however, we still observe enrichment of IEGs among downregulated DEGs in L4/5 IT neurons when controlling for epilepsy status (Supplementary Fig. 14; Supplementary Data S13). Larger, deeply phenotyped cohorts and temporally resolved systems will be required to clarify causality.

Together, our findings suggest that altered transcription factor activity, particularly involving RFX family genes and associated downstream regulatory programs, may contribute to cell-type-specific vulnerability in BA22.

Linking these molecular alterations to language phenotypes provides a framework for understanding how disrupted regulatory architecture in defined neuronal populations could influence verbal ability in ASD^11^. This work represents a step toward integrating chromatin state, transcriptional regulation, and behavioral heterogeneity within a single regional context in NDD/ASD.

## Methods

Code for all quality control and analysis can be found here: https://github.com/sanderslab/brainnet-public.git.

### Subjects/ABN

Brodmann Area 22 (BA22) tissue punches were obtained from donors through the Autism Brain Net (ABN; *N* = 86) and the NIH NeuroBioBank (*N* = 15 individuals). Donor metadata, including postmortem interval, sex, age, hemisphere, intellectual functioning, verbal status, and additional clinical diagnoses, were used to define a cohort with broadly comparable characteristics (Supplementary Data S1). Clinical information on intellectual functioning and verbal status was available for a subset of donors (approximately 20 and 26 individuals, respectively) at the time of tissue acquisition from the biobank.

### Nuclei Isolation and sn-multiome generation

Experimenters isolating nuclei from all donors provided by Autism Brain Net were blinded to sample genotype, age, and sex. All donor tissue samples were shipped with unique shipping IDs from Autism Brain Net or the NIH and stored until nuclei or DNA extraction for SNP genotyping. Frozen tissue blocks from individuals were dissected and cut in groups of 10-12 individuals on dry ice, specifically targeting grey matter. An equal amount of each sample was obtained using this method and added to achieve a final weight of 50-100mg of pooled tissue in a shared tube of frozen tissue. The resulting tube contained a pool of frozen tissue with equal weight representation from each sample. The tube containing the pool of frozen tissue was then homogenized using a pre-chilled Dounce and a homogenization buffer consisting of 0.2653 M Sucrose, 26.6 mM KCl, 5.31 mM MgCl_2_, 21.2 mM Tricine-KOH, pH 7.8, Ultrapure H_2_O, 1 mM DTT, 0.5 mM Spermidine, 0.15 mM Spermine, 0.3% NP40, 1X Protease Inhibitor, and 0.6 U/uL RNAse Inhibitor. The resulting homogenized mixture was then centrifuged to pellet the sample, and the supernatant was removed. The nuclei within the sample were isolated using an iodixanol gradient, through which varying iodixanol concentrations separated the nuclei band from the debris and myelin. The myelin and debris, resting at the top of the gradient, were extracted and discarded to ensure clean nuclei recovery. The nuclear band was extracted from the gradient and diluted with wash buffer containing 10 mM Tris-HCl, pH 7.4, 10 mM NaCl, 3 mM MgCl2, 1% BSA, 0.1% Tween-20, 1 mM DTT, 0.6 U/uL RNAse Inhibitor, and ultrapure H2O. The resulting diluted sample was then counted manually using a hemocytometer and Trypan Blue staining. The solution was then centrifuged, and the nuclear pellet was resuspended to a concentration of 10,000-12,000 nuclei/uL. The final nuclei were suspended in the recommended Diluted Nuclei Buffer containing 1X Nuclei Buffer (PN-2000207, 10x Genomics), 1 mM DTT, 1 U/uL RNase Inhibitor, and Nuclease-Free H2O as recommended. The suspended nuclei resulting from the nuclei isolation protocol were captured at least 4 times in 10x Genomics Multiome wells with around 40,000 nuclei input per well. The target nuclei recovery was about 18,000-20,000 nuclei per well. The following steps were followed per the Multiome protocol using 10x Genomics’ Chromium Next GEM Single Cell Multiome ATAC + Gene Expression User Guide Rev E. The resulting Gene Expression and ATAC libraries underwent quality control and quantification using a High Sensitivity DNA Bioanalyzer 2100 (Agilent) to ensure adequate concentration and curve quality before sequencing. Equal moral ratios of all libraries were sequenced through the UCSF Center for Advanced Technology (CAT) and the Fabrication and Design Center. The Gene Expression libraries were sequenced on a Novaseq X 10B/25B flow cell under the following parameters: Read 1 of 28 bp, Index 1 of 10 bp, Index 2 of 10 bp, and Read 2 of 90 bp. The targeted reads per cell for the Gene Expression libraries were 25,000 reads. The ATAC libraries were sequenced on a Novaseq X 25B flow cell under the following parameters: Read 1 of 151 bp, Index 1 of 12 bp, Index 2 of 24 bp, and Read 2 of 151 bp.

ATAC libraries were targeted for 30,000 reads. All sequenced libraries utilized the recommended 1% PhiX spike-in. All libraries were sequenced one or more times to achieve a final satisfactory read depth (see Supplementary Data S2).

### DNA extraction for SNP array

Donor tissues were cut on dry ice to obtain a 20mg aliquot and stored in -80 °C until DNA extraction. DNA Extraction for SNP genotyping assay was performed using the PureLink Genomic DNA extraction kit (CAT: K182001) using protocols from the manufacturer. Sample quality and concentrations were measured on a NanoDrop One spectrophotometer. All samples that passed QC were > 1.6-1.9 for 260/280 and contained a total of > 2000 ng of DNA. Isolated DNA from samples was sequenced using the iSCAN instrument (Illumina) for the Global Screening Array (GSAv2) chip at the QB3 facility at the University of California, Berkeley.

### Genotype quality control (QC) and imputation

The resultant genotype array data (GSAv2, GRCh38) were filtered for quality control and imputed to the TOPMED imputation panel^65^. Genotype QC was conducted using PLINK v1.90 b7 (16 Jan 2023). Samples were checked for discordant sex (1 sample was ambiguous), high heterozygosity (± 3 SD from the mean), high missingness (± 3 SD from the mean), and relatedness. One sample failed all QC measures, and an additional three were found to have high heterozygosity; however, this was explained by having “admixed” ancestry or ancestry divergent from the majority of samples through principal component analysis (PCA) done with 1,000 Genomes^66^. Two samples were identified as identical via identity by descent analysis (Z0 < 0.05 and Z1 < 0.05). There was no indication in the phenotype data that these two samples were twins (different age, case status, and cause of death), suggesting a sample mix-up. These were removed. SNPs were filtered for high call rate (≥ 0.98) and Hardy-Weinberg Equilibrium (*P* < 1x10^-6^). Ambiguous (aka palindromic A/T, G/C) SNPs and monomorphic SNPs (i.e., SNPs where everyone is homozygous) were removed. SNPs were prepared for imputation against TOPMED using a check tool by Will Rayner and can be found here: https://www.chg.ox.ac.uk/∼wrayner/tools/. The script compares strand, alleles, position, reference, and alternate (REF/ALT) assignments and frequency differences between the data and the reference panel. It subsequently updates strand, position, and REF/ALT assignment to match the reference and removes ambiguous SNPs if MAF > 0.4, SNPs with differing alleles, SNPs with > 0.2 allele frequency difference, and SNPs not in the reference panel. Post-imputation, SNPs were filtered for MAF ≥ 5% and imputation R^2^ ≥ 0.95.

### Principal Component Analysis (PCA) of genotype array data

Principal components were generated through PCA using PLINK v2.00a4.2LM (31 May 2023). QC-ed SNPs were a subset of those overlapping with unrelated individuals in 1,000 Genomes (up to 2nd-degree relatives were removed). We used linkage disequilibrium (LD)-pruned SNPs (pairwise r^2^ <0.2 in batches of 50 SNPs with sliding windows of 5) with MAF > 5% and removed 24 regions with high or long-range LD, including the HLA, leaving 294,312 variants for PCA.

### Read preprocessing, alignment, and count generation

Read preprocessing, alignment, and count generation were done using 10x CellRanger-ARC version 2.0.2 with genome reference 2020-A (July 7, 2020), which uses GRCh38 primary assembly from Ensembl 98 and gene annotations/structure from GENCODE v32 primary assembly. CellRanger-ARC was run per 10x well to identify well-specific quality control metrics.

### Ambient RNA detection

We used CellBender v0.3.0 with default settings on unfiltered CellRanger ARC matrices to identify barcodes likely tagging ambient RNA rather than nuclei. Barcodes marked as ambient RNA were removed. Downstream QC and analysis were done on unfiltered matrices from CellRanger-ARC, with barcodes identified as ambient RNA by CellBender removed.

### Donor assignment

Donor assignment of barcodes was done using demuxlet with modified code from Vincent Gardieux (https://github.com/VincentGardeux/demuxlet.git) to avoid issues with random access memory (RAM) when using a large number of SNPs. As input to demuxlet, we included QC-ed and imputed genotype data (See “Genotype quality control and imputation”) and bam files output from 10x CellRanger-ARC version 2.0.2 (See “Read preprocessing, alignment and count generation”). Three donors had failed genotype array QC. Consequently, we used QC-ed WGS data, filtered to variants matching QC-ed genotype array data, to demultiplex these three donors.

### Identity matching across data modalities

To ensure donor identities were consistent across genotype array data, whole-genome sequencing (WGS), metadata, and multiome data, we performed two complementary identity checks: (1) identity-by-descent (IBD) analysis between WGS and genotype array data, and (2) sex concordance checks across metadata, genotypic data, and expression data (based on expression of *XIST* and Y-chromosome genes).

First, we performed IBD analysis to detect potential sample swaps between genotype array data (used for demultiplexing) and WGS data (used to identify putatively damaging variants associated with ASD and other neurodevelopmental disorders). This analysis revealed that two donors in the genotype array dataset matched each other in the WGS dataset, suggesting a potential sample misassignment.

We next examined sex concordance across datasets. For these two donors, the metadata-reported sex matched the genotypic sex inferred from the WGS data but was inconsistent with the genotypic sex inferred from the genotype array data. In addition, expression patterns of sex-linked genes in the multiome data were inconsistent with the metadata assignments: nuclei assigned to these donors showed expression of Y-chromosome genes (e.g., *UTY* and *ZFY*) with little or no *XIST* expression, or vice versa, contrary to the expected patterns based on the metadata.

Taken together, these results strongly suggested that the labels for these two donors had been swapped in the genotype array dataset. We therefore reassigned the nuclei associated with these donors accordingly.

IBD analysis also identified an additional mismatch between the genotype array and WGS datasets, in which a donor in the genotype array data matched a different donor in the WGS data. Because WGS data were available for only one of these donors, we could not determine whether this represented a reciprocal swap or mislabeling in the genotype array dataset. Additionally, multiome expression for this donor showed low expression of Y-chromosome genes, inconsistent with the male sex indicated in the metadata and genotype. Because the donor identity could not be confidently resolved, this donor was excluded from downstream analyses.

### Cell and gene level quality control

After identifying and removing failed wells, we concatenated RNA count matrices across all wells. Barcodes were first restricted to high-confidence singlets and non-ambient droplets by intersecting demultiplexing assignments run on unfiltered CellRanger ARC matrices (demuxlet; retaining only singlets) and CellBender output (removing ambient RNA-contaminated barcodes, also run on unfiltered CellRanger ARC matrices). We removed genes expressed (count > 0) in ≤ 3 cells. QC filtering was then applied to this subset of barcodes (Supplementary Fig.15-18).

Barcode-level quality control (QC) was performed using combined RNA and ATAC sequencing metrics computed per cell barcode in a Seurat/Signac framework with BPCells-backed matrices^67–69^. For each well, 10x Genomics feature-barcode matrices were loaded and used to construct a joint RNA-ATAC Seurat object. ATAC peaks were restricted to standard chromosomes (1-22, X, Y, and mitochondria) prior to QC. RNA-based QC metrics included total RNA counts, number of detected genes, and percentage of mitochondrial reads. Chromatin accessibility (ATAC) metrics computed per barcode included transcription start site (TSS) enrichment, n ATAC, nucleosome signal, percent reads in peaks, fraction of reads overlapping promoter regions, and fraction of reads overlapping ENCODE blacklist regions (Supplementary Fig. 15-18). Log-transformed versions of selected metrics (n genes, n count RNA, and n count ATAC) were computed using log(1 + x).

Two filtering strategies were used: (i) median absolute deviations (MADs) based filtering and (ii) absolute thresholds. For MAD-based filters, barcodes were flagged as outliers if the metric value lay more than *k* MADs above or below the median of that metric, where *k* is the specified threshold. For absolute thresholds, barcodes above or below the specified threshold(s) were flagged.

The following QC criteria were applied per barcode:

RNA:

- **log(nFeature_RNA + 1)**: MAD-based filtering with k = 4
- **log(nCount_RNA + 1)**: MAD-based filtering with with k = 4
- **Percent mitochondrial reads**: > 8% removed

ATAC:

- **log(nCount_ATAC + 1):** MAD-based filtering with k = 4
- **TSS enrichment**: outside-range filter, retaining barcodes with values between 2 and 15
- **Blacklist ratio**: > 5% removed
- **Nucleosome signal**: MAD-based filtering with k = 5
- **Percent reads in peaks**: MAD-based filtering with k = 5
- **Promoter ratio**: MAD-based filtering with k = 4

### RNA normalization, integration, and clustering

The merged RNA count matrices were processed in Seurat v5^67^. Cells flagged as outliers by predefined QC thresholds were removed before analysis. Counts were normalized using SCTransform, which fits a regularized negative-binomial regression to UMI counts and returns variance-stabilized Pearson residuals. Azimuth v0.5.0^70^ was applied to the SCTransformed data to generate predicted cell-type labels, which were retained as metadata but not used to drive clustering. Batches (wells) were integrated using Seurat’s IntegrateLayers() with CCA-based integration, using the SCT assay as input to produce an integrated low-dimensional representation. A shared nearest-neighbor graph was constructed using the first 30 dimensions of the integrated reduction, and clustering was performed with the Louvain algorithm (FindClusters()) at resolution 0.6, and UMAP was computed on the same integrated dimensions for visualization.

### Cell type assignment

Following normalization, integration, and clustering, as well as cell-type annotation using Azimuth v0.5.0, we used the FindAllMarkers function in Seurat to identify the top 25 expressed genes in each cluster. We compared the weighted expression of these markers to datasets^16,71,72^ using the UCSC browser^73^ tool and determined consensus calls on all clusters. The Azimuth cluster calls were then compared to the manually derived cluster calls to establish a confident cluster identification. All clusters were identified as a single cell type. We either removed or merged clusters <900 cells. Those clusters that showed >90% transcriptomic similarities were merged; among these were L5/6NP, SST, and PVALB clusters, thus identifying 26 neuronal and 12 non-neuronal clusters (Fig. 1c, d).

### Interneuron Subclustering

To define sub-clusters of interneurons in the adult samples (>22 years), we used the iterative clustering package hicat^18^. Briefly, cells were grouped into neighborhoods based on their cell type assignment.

Mitochondrial, ribosomal, sex-specific genes, and genes detected in less than 4 cells were removed from the dataset, after which high-variance gene selection and dimensionality reduction were performed. Jaccard-Louvain clustering was applied recursively until differential expression between sub-clusters was no longer significant, as defined by the “DE score” (sum of -log_10_(adjusted p) over significantly differentially expressed genes) crossing a manually-set threshold. This process was repeated 50 times on 80% samples of each neighborhood, then consensus sub-clusters were formed using the proportion of times that cells clustered together. Sub-clusters were then manually curated by removal or merging if the cluster was dominated by low quality or batch effects between samples. Annotations for each sub-cluster were assigned based on marker genes discovered by differential expression between the sub-cluster and its nearest neighbor sub-cluster, other sub-clusters in the cluster, neighborhood, and globally. To define sub-clusters at earlier developmental stages (<22 years), sub-cluster labels were transferred from adult samples to earlier age groups (young adult, adolescent, late childhood, early childhood) sequentially using reciprocal PCA-based label transfer.

### Cell-type proportion analysis

Cell-type proportion differences were assessed at the donor level. For each donor d, we computed the total number of retained cells (Nd) and the number of cells assigned to cluster c, denoted Ndc. Cluster proportions were defined as: pdc = Ndc / Nd. To stabilize variance across clusters, proportions were transformed using the arcsine–square root transform: ydc = arcsin(sqrt(pdc)), as implemented in the propeller framework of the speckle package^74^. Clusters were tested only if each comparison group contained ≥5 donors with ≥5 cells in that cluster and at least two donors per group overall. For each eligible cluster, we fit a linear model using limma with empirical Bayes moderation to test the effect of case status. Models included donor-level technical covariates: the median number of detected genes per cell (median nFeature_RNA across retained cells) and total sequencing depth (sum of nCount_RNA across retained cells). Covariates were z-scored prior to model fitting. P values were adjusted for multiple testing across clusters within each contrast using the Benjamini–Hochberg false discovery rate (FDR) procedure.

### Age-dependent gene expression modules

For each cell type, donor-level pseudobulk expression matrices were modeled with the formula:

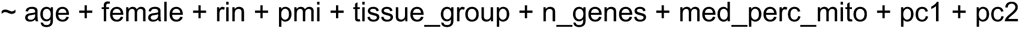

excluding covariates lacking within-group variance. Genes were then ranked by age-VE and assigned to tiers reflecting the strength of their age association. Genes that represented the a, high-confidence were defined adaptively as age-DEGs (Variance explained 0.20, min (Q90(age-VE), 0.25)); and typically, the top ∼10 % per cell types were carried forward for trajectory modeling. For each gene, expression across donors was modeled as a smooth function of age using generalized additive models (GAMs) to capture non-linear developmental trends^75^. Genes with poor or unstable fits (e.g., insufficient deviance explained or over-fitted curves) were excluded. The fitted GAMs were then evaluated on a uniform (0–22) year age grid, generating standardized z-score profiles that summarize each gene’s developmental trajectory. These smoothed gene-wise trajectories were clustered by similarity in shape to identify co-regulated gene groups, referred to as modules. Each module thus represents a distinct temporal pattern of gene expression change across postnatal development, with non-overlapping gene membership and a representative mean trajectory for the cell type.

### Age-DEG trajectories; rate of change and acceleration analysis

To quantify the tempo of developmental transcriptional change, we analyzed the Age-DEG trajectory summaries generated for each cluster. We used the smoothed, z-scored mean expression of Age-DEGs evaluated on an age grid between 0 and 22 years. Within each cluster, we computed the first derivative of the trajectory as the discrete rate of change in z-score between adjacent age points (Δz/Δage) and the second derivative as the change in this rate across age (Δ(rate)/Δage). These derivatives were evaluated at the midpoint of each age interval (age_mid and age_mid_accel). To obtain a robust, cell-type–level summary, we then averaged the absolute value of the rate and acceleration across all Age-DEGs within a cluster, yielding mean |Δz/Δyear| and mean |Δ(rate)/Δyear| as functions of age. We repeated this analysis for all fine clusters that had valid Age-DEG trajectories and annotated each cluster into broad classes (Excitatory, Interneuron, Oligo/OPC, Astro/Microglia, Other) based on its label pattern. For visualization, we generated both class-level overlays and per-cluster curves, and additionally restricted the x-axis to 0–10 years in selected plots to focus on early postnatal dynamics.

### Gene ontology and KEGG analysis (signature modules)

Gene sets for each signature module were compiled from module-specific CSV files containing human gene symbols. Gene names were cleaned, deduplicated, and mapped to Entrez IDs using clusterProfiler and org.Hs.eg.db; modules with fewer than 10 mapped genes were excluded. GO Biological Process (BP) enrichment was performed using enrichGO() with ontology = BP, Benjamini–Hochberg correction, *P* < 0.05, q < 0.20, and a minimum gene set size of 10. KEGG pathway enrichment was performed using enrichKEGG() (organism = *hsa*) with the same significance thresholds. Enrichment was run both per module and on the combined union of all signature genes. All results were exported as CSV files with dot-plot visualizations for significant terms.

### Differential gene expression

RNA counts were pseudobulked by cell type and converted to a SingleCellExperiment object. We used precision-weighted linear mixed models (Dreamlet) to perform differential gene expression (DGE) analysis. Prior to pseudobulking, samples from predefined control-only nuclei preps (batches) were removed, and, for each donor, only nuclei preps with the highest donor cell count were included. Donors with fewer than 300 cells after filtering were excluded. Cells were aggregated to donor x cell-type pseudobulk counts using aggregateToPseudoBulk(), and donor-level covariates (number of cells, total reads, median percent mitochondrial) were computed from the cell metadata and added as covariates. For each cell type we modelled gene counts with dreamlet using a linear model that included case status as the primary contrast and adjusted for age, sex, RNA integrity number (RIN), post-mortem interval (PMI), tissue group, number of genes detected, median percent mitochondrial reads, and the first two genetic principal components (formula: ∼ case status + age + sex + RIN + PMI + nuclei prep + N genes + median percent mitochondrial reads + pc1 + pc2). Covariates that explained more than 1% variance in gene expression were selected. We evaluated pairwise correlations among both donor-level and cell-level covariates and excluded covariates with Pearson correlation (r²) ≥ 0.6 to avoid collinearity. For cell-type-specific covariates, exclusion was applied if this threshold was exceeded in any cell type. As there was only one female individual among individuals without a genetic diagnosis, DGE analysis was conducted among only male cases and controls for the ASD comparison. Genes were filtered by default dreamlet settings (min.cells = 5, min.count = 5, min.samples = 4, min.prop = 0.4). To check for inflation of results, we ran differential expression analysis with shuffled case status labels, and plotted quantile-quantile (QQ) plots of the resultant p values per cell type (Supplementary Fig. 19-21).

### Differential accessibility analysis

Pseudobulked chromatin accessibility data were analyzed using a Seurat/Signac framework with BPCells-backed matrices. Donor-level pseudobulk samples were annotated with covariates including age, sex, RIN, PMI, nuclei prep, total peak fragments, transcription start-site (TSS) enrichment, and the first two genetic principal components. Analyses were performed separately for each cell type. For each cell type, peaks were filtered to retain those with counts per million (CPM) > 1 in at least 40% of donors. Differential accessibility between cases and controls was assessed using DESeq2 likelihood ratio tests (LRT), comparing a full model including case status (∼ case status + age + sex + RIN + PMI + nuclei prep + N fragments + TSS enrichment + pc1 + pc2) against a reduced model excluding case status. Cell types with fewer than 10 valid donor samples were excluded. Multiple testing correction was performed using the Benjamini-Hochberg method.

### Gene-set enrichment analysis (fGSEA)

Differential expression results were analyzed separately for each cell type and comparison group (with genetic diagnosis, without genetic diagnosis, and all). For each cell type, genes were ranked by signed log_2_ fold change, without pre-filtering on statistical significance. Disorder-associated gene sets were derived from MAGMA gene-based association results for common traits (top 500 genes with nominal *P* < 0.05, ranked by significance and SNP count), or from curated rare-variant gene lists for autism and neurodevelopmental conditions. Gene set enrichment analysis was performed using the fGSEA package with 15-500 genes per set, generating normalised enrichment scores (NES), nominal *P* values, and Benjamini-Hochberg-adjusted p values.

### Gene ontology and KEGG analysis (DEGs)

Functional enrichment analyses were performed using the clusterProfiler framework (v4.x) with annotations from org.Hs.eg.db. Differentially expressed genes (DEGs) were identified separately for each cell type and comparison group (without genetic diagnosis, with genetic diagnosis, and all). Genes were considered significant if they met an FDR-adjusted p-value threshold of < 0.05, with no minimum log2 fold-change threshold. For direction-specific analyses, DEGs were further stratified into upregulated and downregulated sets based on the sign of the log2 fold change.

Gene symbols were converted to Entrez Gene identifiers using bitr() before enrichment analysis. Gene Ontology (GO) enrichment was performed for Biological Process (BP), Molecular Function (MF), Cellular Component (CC), and the combined ontology (“ALL”) using enrichGO(), with Benjamini-Hochberg correction applied to all enrichment tests. KEGG pathway enrichment was performed using enrichKEGG() with the organism set to *Homo sapiens*. For both GO and KEGG analyses, terms were considered for visualization if they met a nominal *P-value* cutoff of 0.05, a *q*-value cutoff of 0.2, and contained at least three genes from the input gene set (Supplementary Data S7).

### GWAS Summary Statistics Processing

To evaluate the genetic enrichment of complex traits, GWAS summary statistics were obtained for 25 traits and diseases: age of walking, Alzheimer’s disease, amyotrophic lateral sclerosis, anorexia nervosa, attention deficit hyperactivity disorder, autism spectrum disorder, bipolar disorder, childhood epilepsy, developmental disorders, educational attainment, epilepsy, frontotemporal lobar degeneration, generalized epilepsy, height, intelligence, major depressive disorder, multiple sclerosis, obsessive-compulsive disorder, panic disorder, Parkinson’s disease, post-traumatic stress disorder, problematic alcohol use, schizophrenia, substance use disorder, and Tourette’s syndrome. We harmonized the summary statistics using MungeSumstats (v1.14.1)^76^. Quality control during harmonization included the removal of indels and the exclusion of SNPs with an imputation information score below 0.7, where available.

Gene-level association analyses were performed using MAGMA (v1.08) ^77^, as implemented in FUMA (v1.5.2)^78^. SNPs were assigned to protein-coding genes using a ±1 kb window upstream and downstream of each gene boundary. MAGMA was used to test each gene containing assigned SNPs, and the resulting protein-coding genes were subsequently ranked by their MAGMA P-values to identify the top trait-associated genes.

### Stratified LD Score Regression and Enrichment Analysis

To estimate trait enrichment within specific regulatory regions, we employed linkage disequilibrium score regression (LDSC v2.0.1)^79^. Before LDSC analysis, cell-type-specific peaks were lifted over from the hg38 to the hg19 reference genome using the UCSC liftOver tool (chain file: https://hgdownload.soe.ucsc.edu/goldenPath/hg38/liftOver/hg38ToHg19.over.chain.gz). The lifted-over peaks were formatted for analysis using the make_annotation.py script. Linkage disequilibrium (LD) scores were then computed for each annotation set using ldsc.py. Finally, to rigorously account for multiple testing across traits and annotations, a false discovery rate (FDR) correction was applied to all resulting enrichment p-values.

### Motif activity

Transcription factor motif activity was quantified from scATAC-seq from 10X multiome data using Signac and chromVAR in R^80^. Position frequency matrices from the JASPAR 2024 CORE vertebrate collection were retrieved using the JASPAR2024 and TFBSTools packages ^81,82^. Motif matches within accessible chromatin peaks were identified using Signac’s AddMotifs function with the human reference genome (GRCh38/hg38; BSgenome.Hsapiens.UCSC.hg38).

Motif activity was computed using chromVAR deviation analysis implemented via RunChromVAR. Accessibility counts were aggregated at the pseudobulk level (donors + cell type) before analysis. chromVAR calculates bias-corrected deviation scores by comparing observed accessibility across motif-containing peaks to an expected background matched for technical features such as GC content. Resulting deviation scores were used as quantitative measures of motif activity in downstream analyses.

### Differential motif accessibility

For each cell type, differential motif activity between cases and controls was assessed using linear models implemented in limma, adjusting for age, sex, RNA integrity number, postmortem interval, tissue group, peak fragment counts, and the first two genetic principal components. Donors with missing covariate data were excluded, and case status was modeled as a binary factor. Donors derived exclusively from control-only tissue groups (batches) were excluded prior to analysis. Empirical Bayes moderation was applied, and statistical significance was assessed using Benjamini-Hochberg false discovery rate correction.

### TF activity

We applied the SCENIC+^83^ workflow (v1.0a1) to construct gene regulatory networks (GRNs) from single-nucleus multiomic data. Because the full dataset contained 498,447 nuclei, which is computationally demanding for GRN reconstruction, we subsampled one-tenth of all nuclei while retaining at least 100 cells from each sub-cell type to preserve transcriptional diversity. We next performed topic modeling using pycisTopic, which decomposes the accessibility matrix into latent “topics”, sets of co-accessible regions that may correspond to underlying regulatory programs. The optimal topic number (40) was determined by maximizing the model log-likelihood. To define candidate enhancer regions, three complementary strategies were used together: (1) binarization of topic weights using the Otsu method; (2) selection of the top 3,000 regions per topic; an (3) identification of differentially accessible peaks (log_2_(FC]) > 0.5, adjusted P < 0.05) using the Wilcoxon rank-sum test on imputed accessibility values. These candidate enhancers were subjected to motif enrichment analysis using pycistarget and a discrete element method (DEM)-based approach to identify overrepresented TF-binding motifs, thereby assigning putative TF–region relationships. A custom cisTarget database was built for motif annotation on the 1,450,469 consensus peaks by using “create_cistarget_motif_databases.py” with default parameters: --bgpadding 1000. TF–region– gene regulation network was inferred by using the SCENIC+ snakemake pipeline with default parameters. eRegulons were then defined as TF–region–gene triplets, comprising a TF, all regions enriched for its motif, and the genes linked to those regions based on co-accessibility and correlation patterns. For each eRegulon, specificity scores were computed using the Regulon Specificity Score (RSS) algorithm, based on Area Under the Curve (AUC) enrichment values for either region- or gene-based eRegulons. The top eRegulons per cell type were visualized as cell-type–specific regulatory modules. We extended our eRegulon enrichment analysis from the 49,454 sketched nuclei to all 498,447 nuclei by computing the gene-based AUC scores for all 385 (direct) eRegulons using the R package AUCell (v.1.20.2) using the default settings^83–85^.

### Differential TF activity analysis

As in the differential motif accessibility analysis, pseudobulked chromatin accessibility data were analyzed using a Seurat/Signac framework with BPCells-backed matrices. Direct regulon (eRegulon) activity scores from SCENIC+ were extracted for each donor and analysed separately within each cell type. Donors derived exclusively from control-only tissue groups (batches) were excluded prior to analysis. For each cell type, differential TF activity between cases and controls was assessed using linear models implemented in limma, adjusting for age, sex, RNA integrity number, postmortem interval, tissue group (batch), peak fragment counts, and the first two genetic principal components. Donors with missing covariate data were excluded, and case status was modeled as a binary factor. Empirical Bayes moderation was applied to improve variance estimation, and significance was assessed using Benjamini-Hochberg false discovery rate (FDR) correction.

### Correlation of DEG logFC for autism with and without a diagnosis

To evaluate the concordance between genetic diagnosis and non-genetic diagnosis differential-expression signatures, we performed a repeated random-split analysis (n = 1,000). Samples from predefined control-only tissue groups were removed, and, for each donor, only cells from the donor’s most-represented tissue were retained; donors younger than 10 years (to avoid random sampling from largely mismatched age groups between cases and controls) or with fewer than 300 cells (after filtering) were excluded. Genes on sex chromosomes were removed before analysis (likewise to avoid discordant results due to sex imbalance between cases and controls). For each random iteration, control donors were randomly partitioned into two non-overlapping sets, and two case-control comparisons were constructed: (i) a “with genetic diagnosis” comparison combining donors with known risk loci against control set A, and (ii) a “without genetic diagnosis” comparison of autism cases against control set B. For each comparison and for every cell type, cells were aggregated to donor × cell-type pseudobulk counts, and differential expression was tested using the dreamlet pipeline with the model ∼ case_status + age + sex + RIN + PMI + nuclei prep + N genes + median percent mitochondrial reads + pc1 + pc2. Per-cell-type log_2_ fold-changes for the case status coefficient from the with and without genetic diagnosis contrasts were paired by gene, and Spearman correlation of logFC values was computed within each cell type. Correlations from all splits were recorded and summarized per cell type by the median and 95% bootstrap (without replacement) percentile interval (2.5th-97.5th percentiles).

### Enrichment of DEGs for autism and NDC genes

To assess enrichment of autism and NDC genes among DEGs, Fisher’s exact tests were performed using autism (185) and NDC genes (664) from ^3^. For each comparison (all DEGs and DEGs from donors with a genetic diagnosis), genes were classified as significant if they showed FDR-adjusted *P* < 0.05 in pseudobulk differential expression analyses across cell types. The background universe was defined as all genes tested in the corresponding differential expression analysis. For each gene set, 2x2 contingency tables were constructed contrasting DEG versus non-DEG status and gene set membership, and odds ratios, 95% confidence intervals, and two-sided *P* values were computed using Fisher’s exact test.

### Identification of variants in WGS data

Whole-genome sequencing (WGS) data were generated from DNA extracted from Brodmann Area 10 (BA10). Data generation was performed at the New York Genome Center (NYGC) in twelve batches (WGS1 to WGS12). Sequencing libraries were prepared using KAPA Hyper PCR+ and paired end sequencing was performed using 150 bp read lengths on the Illumina NovaSeq 6000 (WGS1-WGS10) or Illumina NovaSeq X (WGS11-WGS12). Data were processed aligned to GRCh38 using the functional equivalence GATK pipeline ^86^ using BWA-mem and Picard MarkDuplicates. Individual gVCFs were generated with the GATK pipeline using ApplyRecalibration, GenotypeGVCFs, HaplotypeCaller, and VariantFiltration. Variants were annotated using VEP (v104) using ENSEMBL/Gencode (v38) gene definitions, gnomad (v3), ClinVar (20210102), HGMD public (20204), dbSNP (v154), PolyPhen (v2.2.2), and SIFT (v5.2.2). High and moderate effect variants (e.g., PTVs and missense) and Likely Pathogenic and Pathogenic variants in ClinVar were extracted. Joint genotyping was also performed with GATK to increase sensitivity. Structural variants were detected using Manta, Canvas, and the SV-GATK pipeline ^87^. WGS data were available for 37 of the 49 cases and all 37 of the Autism BrainNet controls (WGS data were not available for the NIH controls). Variants detected by WGS were compared with clinical genetics reports from donors. No variants with clear contribution to ASD or NDD were detected in the controls. In the cases, eight copy number variants and five single gene disorders were validated. All variants are indicated in the donor-level metadata (Supplementary Data S1).

### Enrichment of differential eRegulon target peaks in functional genomic regions

To identify regulatory differences between verbal and nonverbal individuals with ASD, we used SCENIC+ to infer differential eRegulons from the 10x multiome data within each annotated cell type. For each differential eRegulon from an individual cell type, we extracted its directly regulated target peaks, defined by SCENIC+ peak-to-gene linkages and transcription factor motif support.

We next evaluated the enrichment of these eRegulon target peaks within specific evolutionary and functional genomic elements. These annotations included human accelerated regions (HARs), human ancestor quickly evolved regions (HAQERs), fetal brain human-gained enhancers (HGEs), primate-conserved regions, and matched random control regions. Enrichment was assessed using Fisher’s exact tests to compare overlap frequencies between the eRegulon target peaks and the annotated genomic regions. For each cell-type-specific test, the background signal was restricted to all accessible ATAC peaks detected in that corresponding cell type. Odds ratios and associated *P*-values were calculated to quantitatively assess the enriched association of differential eRegulon target peaks with these lineage-specific and conserved regulatory elements.

## Acknowledgements

We would like to thank Dr. Alan Packer and Dr. Marta Benedetti of the Simons Foundation, Kelly Gleason (UT Southwestern), Dr. David Amaral and team at UC Davis, and Autism BrainNet for their efforts. Tissues and data were obtained from Autism BrainNet, a resource of the Simons Foundation Autism Research Initiative (SFARI). Autism BrainNet also manages the Autism Tissue Program (ATP) collection, previously funded by Autism Speaks. We are grateful and indebted to the families who donated tissue for research purposes to Autism BrainNet and the ATP.

This project was supported by grants from the National Institutes of Health: R01MH122681, R01MH125516, R01MH128364, R01NS123263, R01MH129751, UM1MH130991, 1S10OD028511-01. This project was also supported by the Simons Foundation Autism Research Initiative Sex Differences Collaboration (SFARI #736613), as well as gifts from Esther A. & Joseph Klingenstein Fund, the Shurl and Kay Curci Foundation, the Sontag Foundation, William K. Bowes Jr. Foundation. T.J.N. is a New York Stem Cell Foundation Robertson Neuroscience Investigator. The UCSF CAT is supported by UCSF PBBR, RRP IMIA.

## Contributions

Conceptualization: V.S., E.M.W., Y.H., S.J.S., T.J.N.

Methodology: V.S., E.M.W., R.L., C.E.

Formal analysis: V.S., E.M.W., Y.H., J.A., M.G.

Investigation: V.S., E.M.W., R.L., C.E.

Resources: T.M., S.J.S., T.J.N.

Data Curation: V.S., E.M.W., Y.H., M.G., N.R., J.W., D.S.P., F.H.L., C.D., N.S., S.D., G.Y.

Writing - Original Draft: V.S., E.M.W, Y.H., R.L., S.J.S., T.J.N.

Writing - Review & Editing: V.S., E.M.W., Y.H., R.L.,G.Y., S.J.S., T.J.N.

Visualization: V.S., E.M.W.,Y.H.,

Supervision: S.J.S., T.J.N.

Project administration: V.S., E.M.W.

Funding acquisition: S.J.S., T.J.N.

