## Supplemental data for "Molecular dynamics of Brodmann Area 22 in development and autism"

### Supplementary Data Legends

**Supplementary Data S1.** Metadata for all donors included in the study. The table reports individual donor identifier (donor_id), patient_id, age, sex, diagnosis, Case_group, case_status, RNA integrity number (rin), postmortem interval (pmi), genomic variant annotation (Variant (GRCh38)), variant consequence (effect), associated risk gene or locus (autism_risk_loci), syndrome, genetic_variant_type, epilepsy_status, epilepsy, medical_conditions, psychological_conditions, intellectual_functioning, verbal_status, shipment_id, tissue_group, and genotype principal components (pc1–pc5).

**Supplementary Data S2.** Summary of sequencing composition and quality control metrics for each 10x well. The table reports well identifier (well_name), total number of nuclei recovered (n_cells), number of donors contributing to the well (n_donors), donor identifiers (donors_in_well), case group composition (Case_groups_in_well), diagnostic composition (diagnoses_in_well), median RNA UMI counts (median_nCount_RNA), median detected genes (median_nFeature_RNA), median gene counts (median_genes_RNA), median UMI counts (median_umis_RNA), median mitochondrial read fraction (median_percent_mt), median ATAC counts (median_nCount_ATAC), median accessible features (median_nFeature_ATAC), median ATAC fragments (median_atac_frags), median transcription start site enrichment score (median_TSS_enrich), median fraction of reads in peaks (median_pct_peaks), tissue source (tissue_groups_in_well), and additional donor and tissue identifier (shipment_ids_in_well).

**Supplementary Data S3.** Donor-level sequencing and quality control metrics. Summary of sequencing yield and quality control metrics for each donor post-filteration. The table reports individual donor identifier (donor_id), patient_id, age, sex, diagnosis, Case_group, tissue sources contributing to each donor (tissue_groups), additional donor and tissue identifier (shipment_ids), total number of nuclei recovered (n_cells_total), number of nuclei retained after post cell type call based filtering (n_cells_KEEP), total RNA UMI counts (RNA_total_UMI), median RNA UMI counts per nucleus (RNA_median_nCount), mean RNA UMI counts per nucleus (RNA_mean_nCount), median detected genes per nucleus (RNA_median_nFeature), mean detected genes per nucleus (RNA_mean_nFeature), median mitochondrial read fraction (RNA_median_pct_mt), total ATAC counts (ATAC_total_counts), median ATAC counts per nucleus (ATAC_median_nCount), mean ATAC counts per nucleus (ATAC_mean_nCount), median accessible features per nucleus (ATAC_median_nFeature), mean accessible features per nucleus (ATAC_mean_nFeature), median transcription start site enrichment score (ATAC_median_TSS), median fraction of reads in peaks (ATAC_median_pct_peaks), and median fraction of reads mapping to promoter regions (ATAC_median_promoter_ratio).

**Supplementary Data S4.** Cluster annotation and summary statistics. Table summarizing cluster-level metadata across all cells. The table reports final annotated cluster identity (cluster_label), broad cell class (Cell_Type_Group), total number of nuclei assigned to each cluster (n_cells), number of donors contributing cells to each cluster (n_donors), and top marker genes identified per cluster (marker_genes). Cluster annotations were derived from Azimuth predictions and validated using marker gene expression and reference datasets.

**Supplementary Data S5.** Donor-level cell-type proportion data used for downstream proportion analyses. The table reports individual donor identifier (donor_id), cell-type annotation (cluster_label), number of cells in the cluster per donor (n_cells_cluster), total number of cells per donor (n_cells_total), fraction of cells per cluster per donor (proportion), and analysis group, including control, ASD, and NDD/ASD (group). Proportions are calculated as cluster cells divided by total cells per donor.

**Supplementary Data S6.** Differential gene expression results by cell type and comparison. Differential expression was performed on donor-level pseudobulk RNA data within each annotated cell type using dreamlet. The table reports gene-level statistics, including cell-type annotation (cell_type), HGNC gene name (gene_symbol), log2 fold change (logFC), average expression across samples (AveExpr), moderated t-statistic (t), nominal P value (P.Value), Benjamini–Hochberg adjusted P value (adj.P.Val), log-odds of differential expression (B), standardized z-score (z.std), and analysis group (comparison). Direction of effect is in cases (controls are the reference group).

**Supplementary Data S7.** GO and KEGG enrichment results for signature gene sets. Gene Ontology (GO) and KEGG pathway enrichment analyses were performed using clusterProfiler. The table includes analysis group (comparison), direction of differential gene expression in cases (direction_in_cases), enrichment type, e.g. GO or KEGG (enrichment_type), GO category (ONTOLOGY), term identifier (ID), term name (Description), proportion of input genes in the term (GeneRatio), proportion in background (BgRatio), enrichment ratio (RichFactor), observed vs expected enrichment (FoldEnrichment), directional enrichment score (zScore), nominal P value (pvalue), adjusted P value (p.adjust), FDR q value (qvalue), overlapping genes (geneID), number of overlapping genes (Count), and optional category and subcategory annotations for KEGG enrichment.

**Supplementary Data S8.** fGSEA results for disorder-associated gene sets. Gene set enrichment analysis was performed using fast gene set enrichment analysis (fGSEA) on ranked differential expression signatures for each cell type and comparison. The table reports gene set name (pathway), cell-type annotation (cell type), normalized enrichment score (NES), nominal P-value (p-value), adjusted P-value (p-adjusted), core contributing genes (leading edge), and analysis group (comparison).

**Supplementary Data S9.** Differential accessibility results for ATAC peaks. Pseudobulk chromatin accessibility was analyzed using DESeq2 likelihood ratio tests within each cell type. The table includes cell-type annotation (cell_type), genomic coordinates hg38 of ATAC peak (peak), mean accessibility across samples (baseMean), log2 fold change in accessibility (log2FoldChange), standard error of log2FoldChange (lfcSE), test statistic (stat), nominal P value (pvalue), adjusted P value (padj), and analysis group (comparison). Direction of effect is in cases (controls are the reference group).

**Supplementary Data S10.** Differential motif accessibility results by cell type. Motif activity (chromVAR deviation scores) was tested for differential accessibility between cases and controls using limma. The table includes cell-type annotation (cell_type), motif ID (motif), associated transcription factor (TF_name), log2FC difference in motif activity (logFC), average motif activity (AveExpr), moderated t-statistic (t), nominal P value (P.Value), adjusted P value (adj.P.Val), log-odds of differential activity (B), and analysis group (comparison). Direction of effect is in cases (controls are the reference group).

**Supplementary Data S11.** Cell-type associations with verbal and intellectual disability status. Statistical results testing associations between cell types and clinical phenotypes. The table reports cell-type annotation (celltype), P-value for association with verbal status comparison (verbal_pvalue), and P-value for association with intellectual disability status comparison (ID_pvalue).

**Supplementary Data S12.** Differential eRegulon activity by cell type. Differential eRegulon activity results across cell types. The table reports cell-type annotation (cell type), transcription factor regulatory module (eRegulon), nominal P-value (p_val), log2 fold change in regulon activity (avg_log2FC), and average regulon activity across samples (baseMean). The direction of effect is relative to the Verbal group.

**Supplementary Data S13.** Immediate early gene enrichment in epilepsy by cell type. Fisher's exact tests were used to assess enrichment of immediate early genes (IEGs) among differentially expressed genes controlling for epilepsy status as a covariate within each cell type. The table reports cell-type annotation (cell_type), analysis group (comparison), number of IEGs that are DEGs (IEG_DEG), non-IEG DEGs (nonIEG_DEG), IEGs not differentially expressed (IEG_notDEG), background genes (nonIEG_notDEG), enrichment effect size (odds_ratio), nominal P value (p_value), and Benjamini–Hochberg adjusted P value (p_adj_bh). The universe is all genes tested in the given cell type.

### Supplementary Figures

#### Supplementary Fig. 1

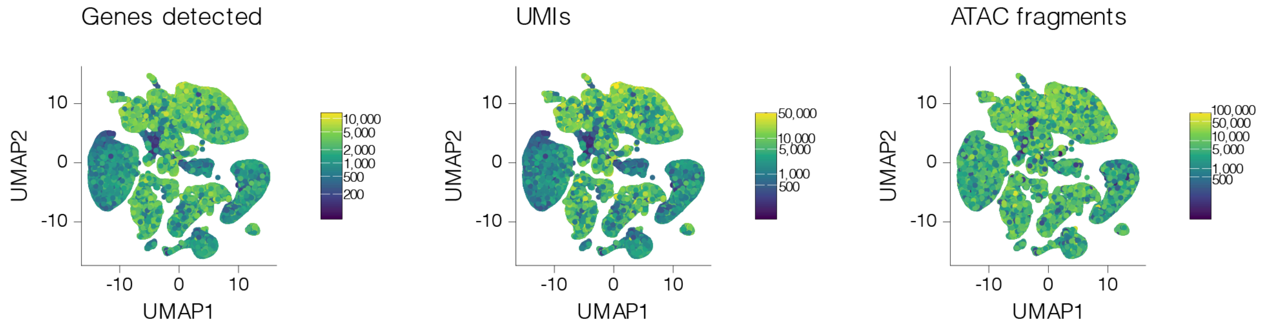

**Supplementary Fig. 1. Library-level quality control metrics.** UMAPs colored by genes detected, UMIs, and ATAC fragments across all cells.

Supplementary Fig. 2

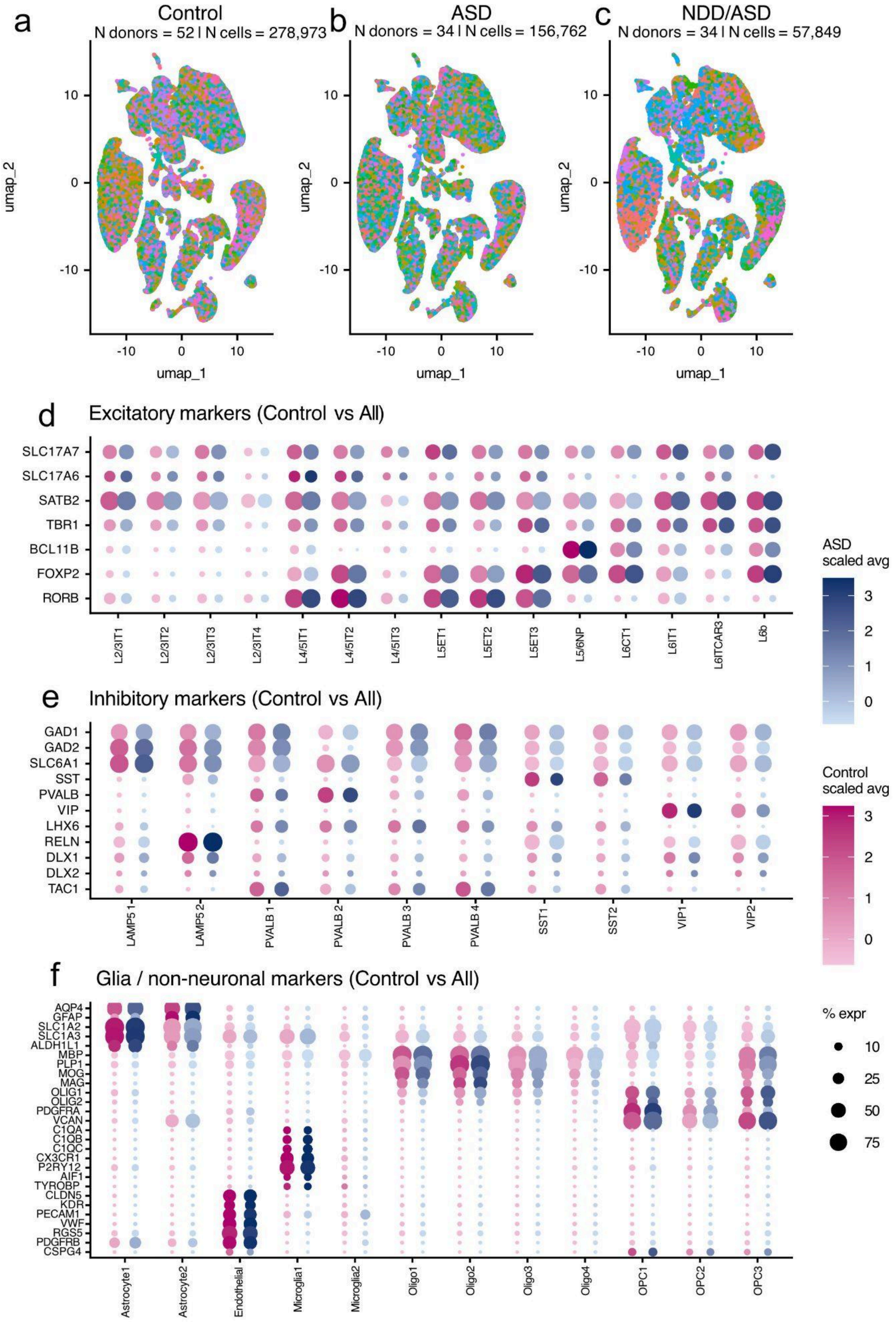

**Supplementary Fig. 2. Cell-type composition and marker expression across diagnostic groups. a-c**, UMAP embeddings of all nuclei colored by donor identity for control donors **a**, ASD without genetic diagnosis (ASD) **b**, and ASD with known genetic diagnosis (NDD/ASD) **c**. For each group, the number of donors and the total number of nuclei plotted are indicated. Identical UMAP coordinates are used across panels to enable direct visual comparison of cellular distributions. **d**, Dot plot showing expression of canonical excitatory neuron markers across excitatory neuronal subclasses. Dot size represents the percentage of cells expressing each gene within a given cluster, and dot color denotes scaled average expression, shown separately for control (magenta scale) and ASD (blue scale) samples. **e**, Dot plot of inhibitory neuron marker expression across inhibitory neuronal subclasses, visualized as in **d**. **f**, Dot plot of glial and non-neuronal marker expression across astrocyte, oligodendrocyte lineage, microglial, endothelial, and perivascular cell populations, visualized as in **d**.

Supplementary Fig. 3

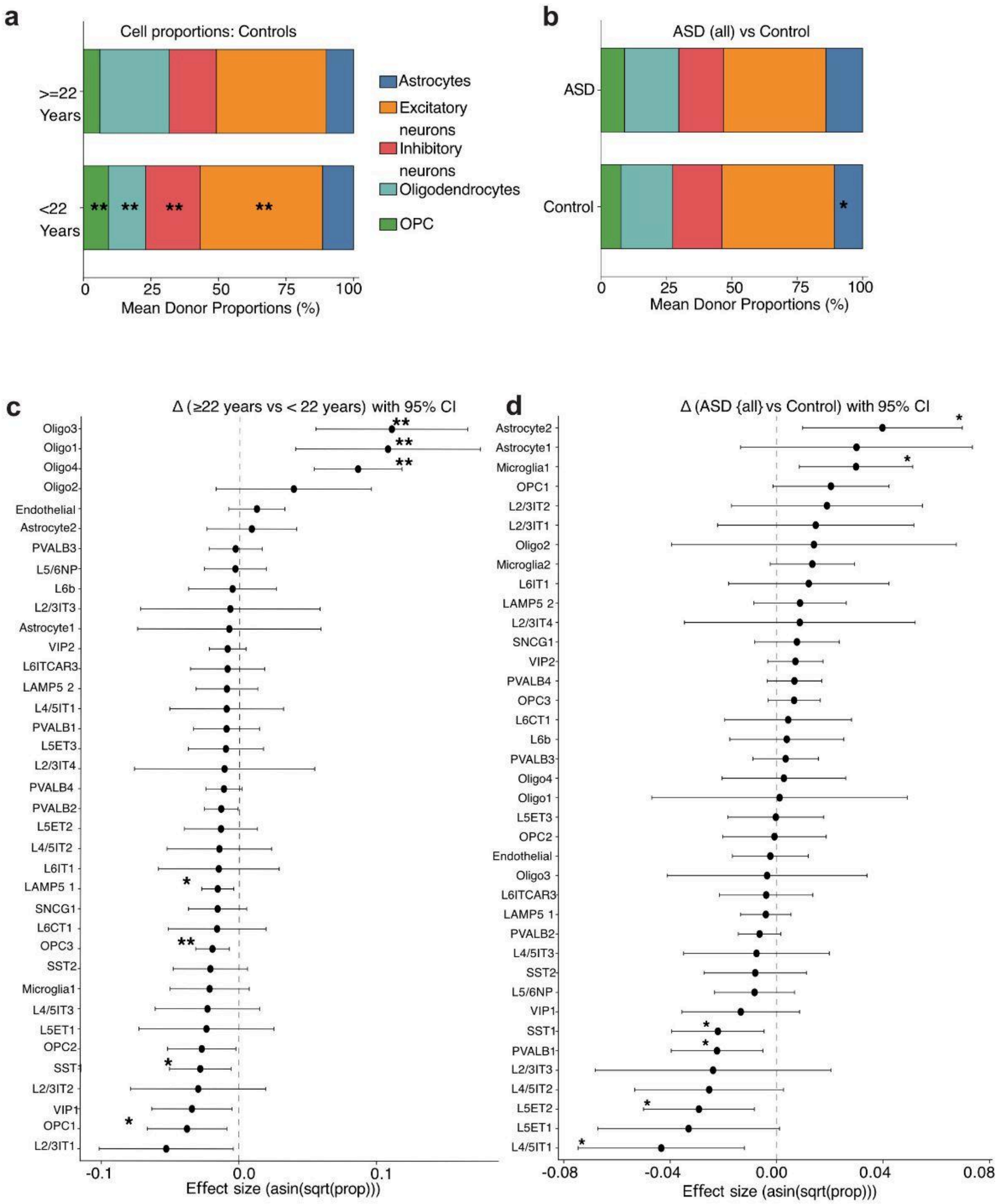

**Supplementary Fig. 3. Changes in cellular composition in BA22. a**, Stacked bar plots showing mean donor proportions of major cell classes in neurotypical individuals stratified by age (<22 years and ≥22 years), illustrating increased oligodendrocytes and decreased OPCs with age. **b**, Stacked bar plots showing mean donor proportions of major cell classes comparing ASD (all) and control individuals, highlighting a modest increase in astrocytes in ASD. **c, d**, Cluster-level effect sizes with 95% confidence intervals for age (Δ ≥22 − <22 years; **c**) and diagnosis (Δ ASD −

Control; **d**, estimated using arcsine-transformed linear models controlling for sequencing depth and gene detection (Methods). (**) indicates FDR < 0.05; (*) indicates FDR < 0.1.

### Supplementary Fig. 4

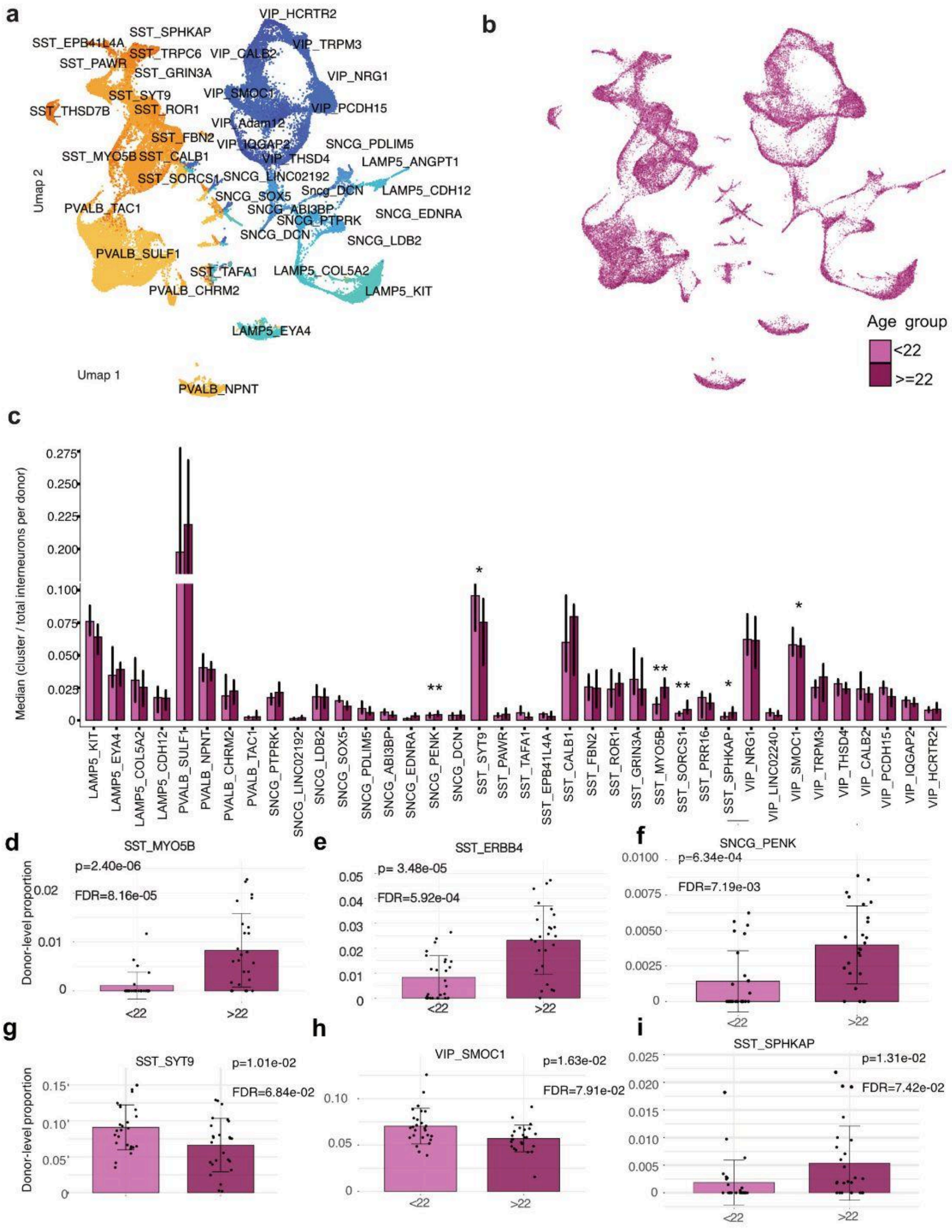

**Supplementary Fig. 4. Developmental dynamics of inhibitory neuronal populations in BA22. a, b**, UMAP embedding of all inhibitory neurons colored by Hi-CAT cluster annotations, **a**, by all Hi-CAT Cluster Annotations, **b**, by age group. **c**, Median cluster proportions of select interneuron classes across donors, showing relative abundance of inhibitory neuron subtypes across the dataset. Bars represent the median proportion of each cluster normalized to total interneurons per donor, with error bars indicating variability across donors, * indicates

FDR-adjusted P-value < 0.1, ** indicates adjusted p-value < 0.05. Statistical significance was estimated using dreamlet linear mixed-effects models with Benjamini–Hochberg FDR correction. **d**-**i**, Donor-level proportions of selected interneuron subtypes showing significant age-associated changes. Each point represents an individual donor, with boxplots summarizing distributions across age groups (<22 years and ≥22 years). Subtypes shown include SST_MYO5B **d**, SST_ERBB4 **e**, SNCG_PENK **f**, SST_SYT9 **g**, VIP_SMOC1 **h**, and SST_SHPKAP **i**.

### Supplementary Fig. 5

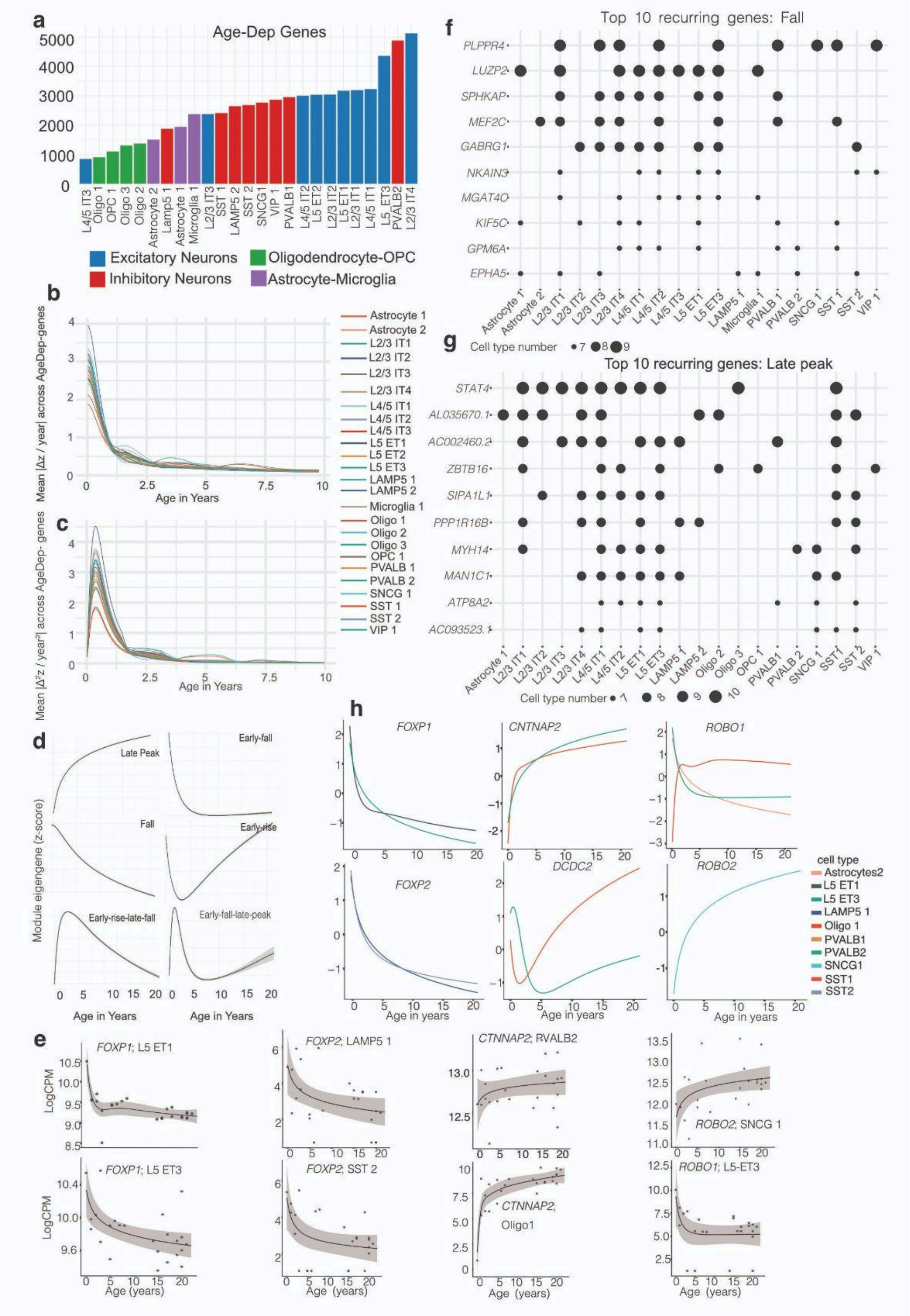

**Supplementary Fig. 5. Postnatal developmental gene programs in human BA22 a,** Number of age-dependent genes (AgeDep genes) identified per cell type using variance partitioning analysis (dreamlet; Methods). Bars are colored by major lineage: excitatory neurons (blue), inhibitory neurons (red), oligodendrocyte–OPC lineage (green), and astrocyte–microglia populations (purple). **b, c,** Mean first **b** and second **c** derivatives of age-associated gene expression trajectories across cell types, illustrating rate and acceleration of transcriptional change during postnatal development. All major lineages exhibit a sharp peak in transcriptional dynamics during early postnatal life, followed by rapid stabilization. **d,** Representative gene expression curves of the six recurrent developmental trajectory modules identified by generalized additive modeling (GAM) and clustering of gene-wise age trajectories. Modules include Early-rise, Early-fall, Early-rise–Late-fall, Early-fall–Late-peak, Late-peak, and Fall patterns (z-scored module eigengenes). **e,** Representative donor-level pseudobulk expression trajectories for selected genes across cell types, illustrating module-specific temporal patterns of language associated genes (e.g., *FOXP1, FOXP2, CNTNAP2, ROBO1, ROBO2, DCDC2*). Lines represent GAM fits; shaded regions indicate 95% confidence intervals. **f, g,** Dot plots showing the top 10 genes most frequently assigned to the Fall **f**, and Late-peak **g**, modules across cell types. Dot size reflects the number of cell types in which the gene exhibits the corresponding trajectory pattern. **h**, Gene expression trajectories for representative genes across selected clusters, highlighting conserved and cell-type–specific developmental dynamics.

### Supplementary Fig. 6

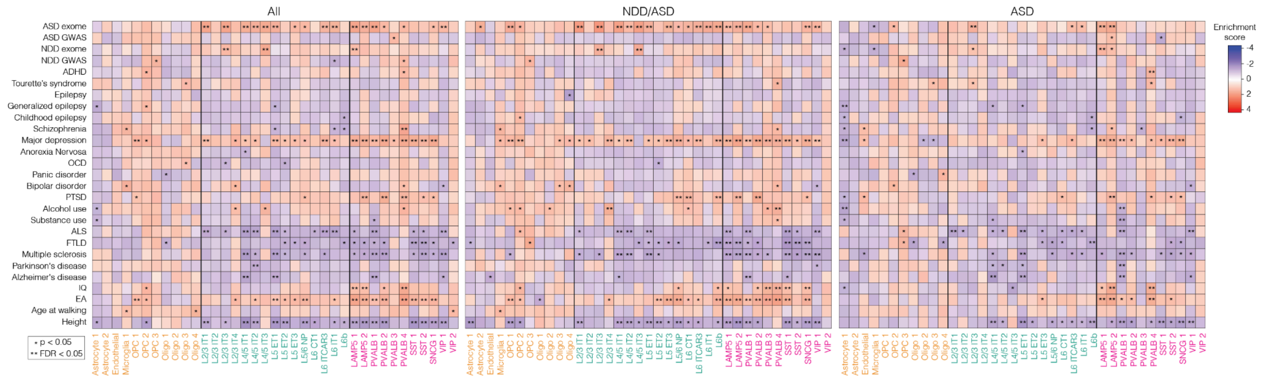

**Supplementary Fig. 6. Gene-set enrichment analysis (fGSEA) by cell type for DEGs from all case vs control comparisons.** Gene sets include the top 500 GWAS genes (Methods) and rare variant gene lists for autism and NDDs. Each heatmap corresponds to a different comparison (All, NDD/ASD, ASD). (Fu et al., 2022; * p < 0.05; ** FDR < 0.05).

### Supplementary Fig. 7

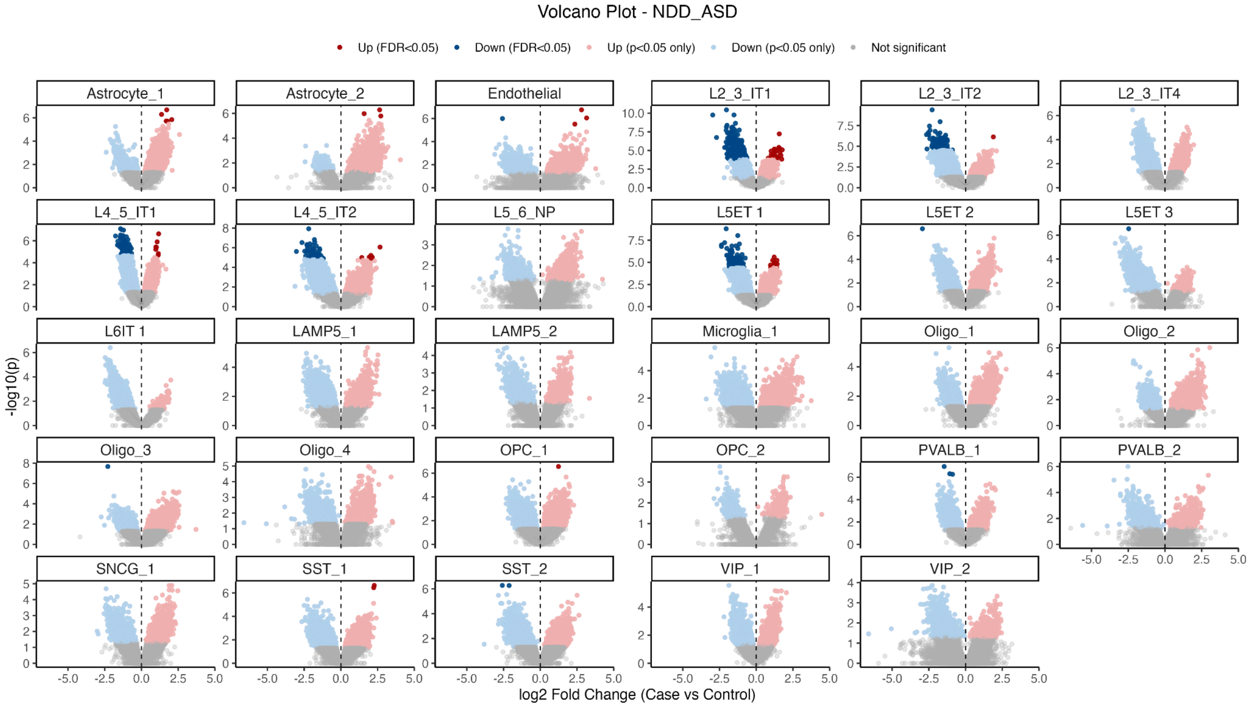

**Supplementary Fig. 7. Cell type specific differential accessibility between NDD/ASD and**

**controls.** Volcano plots showing differentially accessible regions (DARs) across cell types comparing NDD/ASD cases versus controls. Each panel corresponds to a cell type. The x-axis represents $\log_2$ fold change (case vs. control), and the y-axis shows $-\log_{10}$(p-value). Each point represents a peak. Peaks significantly upregulated in NDD/ASD (FDR < 0.05) are shown in red, and significantly downregulated genes (FDR < 0.05) are shown in dark blue. Genes meeting nominal significance only (p < 0.05 but not FDR-significant) are shown in lighter shades (pink for upregulated, light blue for downregulated), while non-significant genes are shown in gray. The vertical dashed line indicates no change ($\log_2$ fold change = 0).

### Supplementary Fig. 8

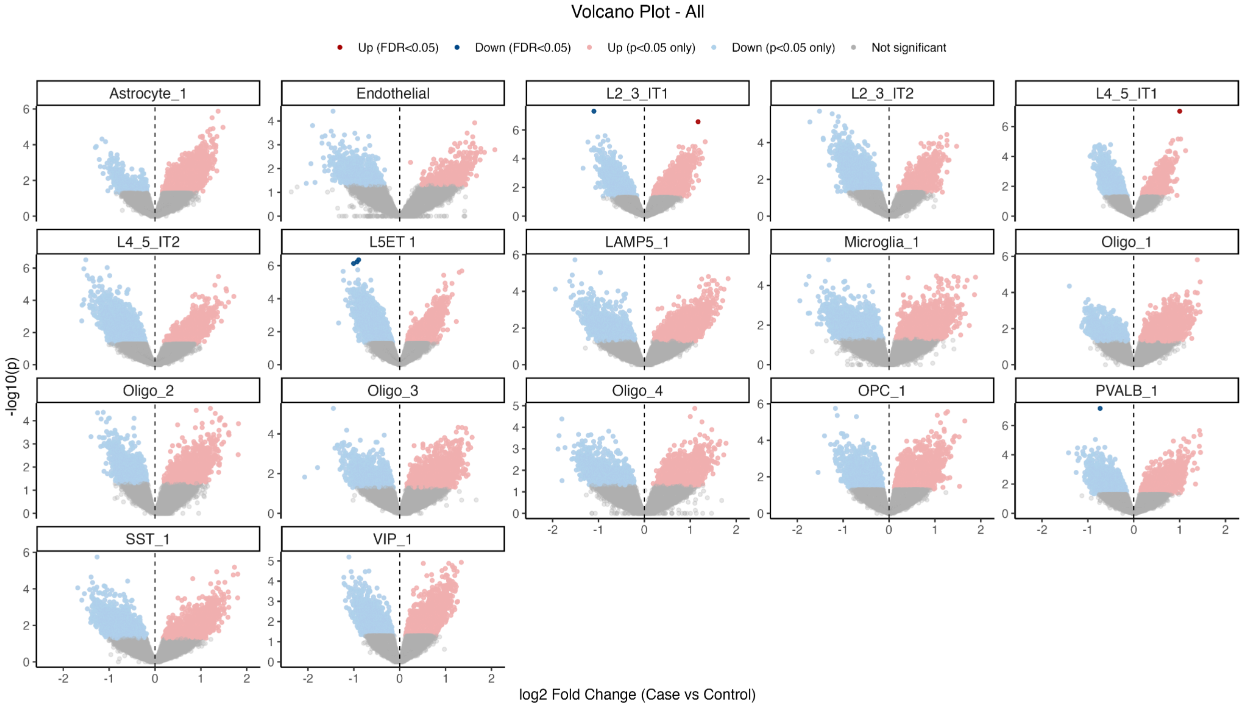

**Supplementary Fig. 8. Cell type specific differential accessibility between all cases and controls.** Volcano plots showing differentially accessible regions (DARs) across cell types, comparing. All cases versus controls. Each panel corresponds to a cell type. The x-axis represents $\log_2$ fold change (case vs. control), and the y-axis shows $-\log_{10}$(p-value). Each point represents a peak. Peaks significantly upregulated in NDD/ASD (FDR < 0.05) are shown in red, and significantly downregulated genes (FDR < 0.05) are shown in dark blue. Genes meeting nominal significance only (p < 0.05 but not FDR-significant) are shown in lighter shades (pink for upregulated, light blue for downregulated), while non-significant genes are shown in gray. The vertical dashed line indicates no change ($\log_2$ fold change = 0).

### Supplementary Fig. 9

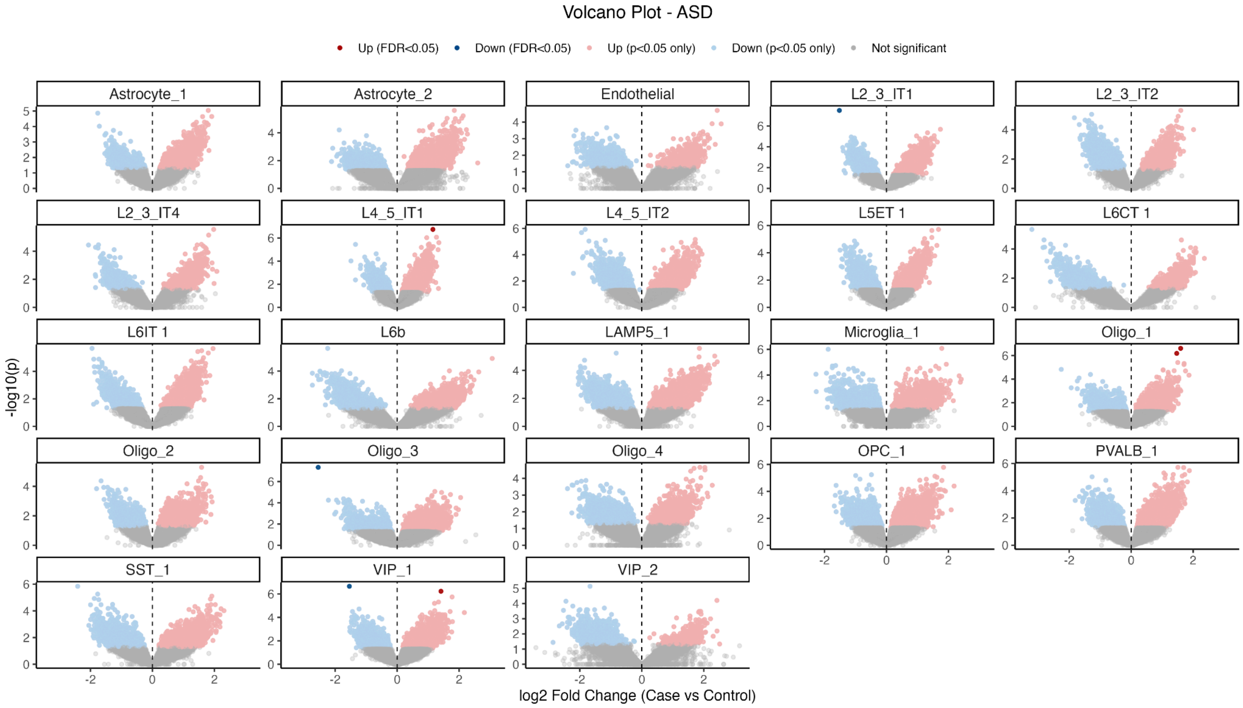

**Supplementary Fig. 9. Cell type specific differential accessibility between ASD cases and controls.** Volcano plots showing differentially accessible regions (DARs) across cell types comparing ASD cases versus controls. Each panel corresponds to a cell type. The x-axis represents $\log_2$ fold change (case vs. control), and the y-axis shows $-\log_{10}$(p-value). Each point represents a peak. Peaks significantly upregulated in NDD/ASD (FDR < 0.05) are shown in red, and significantly downregulated genes (FDR < 0.05) are shown in dark blue. Genes meeting nominal significance only (p < 0.05 but not FDR-significant) are shown in lighter shades (pink for upregulated, light blue for downregulated), while non-significant genes are shown in gray. The vertical dashed line indicates no change ($\log_2$ fold change = 0).

### Supplementary Fig. 10

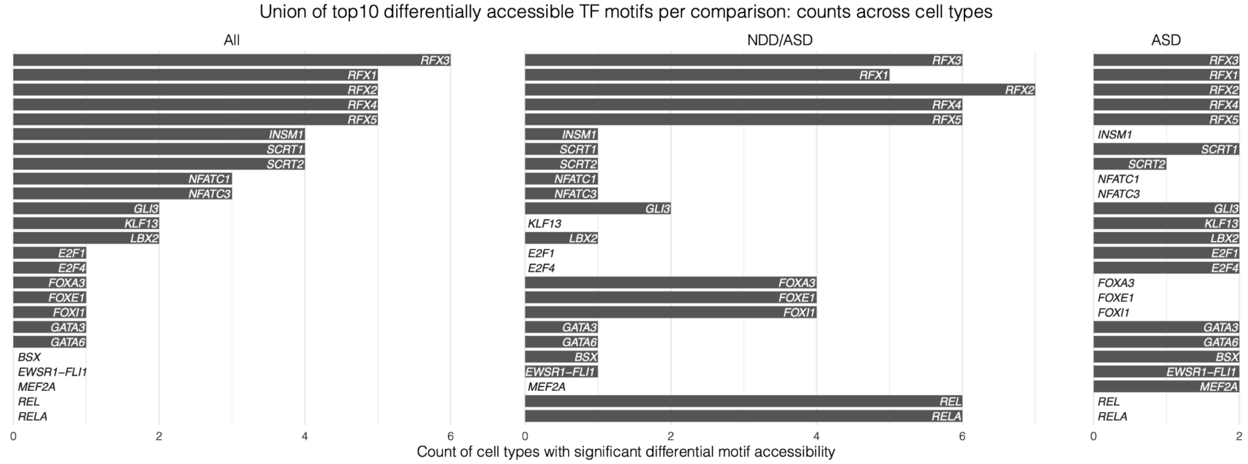

**Supplementary Fig. 10. Union of top differentially accessible transcription factor motifs by case-control comparison.** Bar plots showing the number of cell types in which each transcription factor (TF) motif is significantly differentially accessible (FDR < 0.05). For each comparison (“All”, “NDD/ASD”, and “ASD”), the union of the top 10 most significant TF motifs per cell type was compiled. The x-axis indicates the count of cell types with significant (FDR < 0.05) differential motif accessibility, and each bar corresponds to a TF motif (labeled). Motifs are ordered by decreasing frequency across cell types in the “All” comparison.

### Supplementary Fig. 11

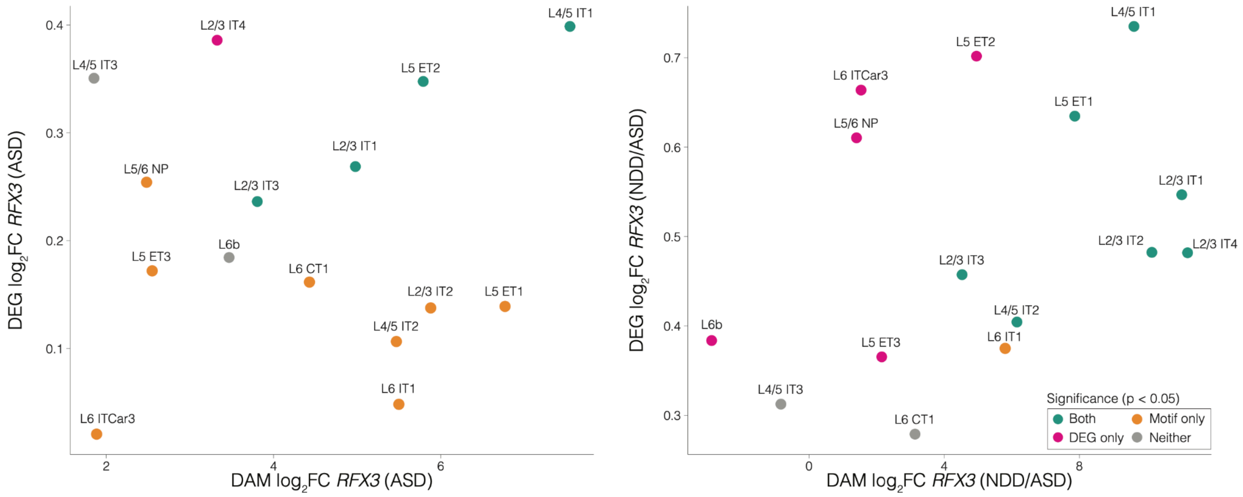

**Supplementary Fig. 11. Relationship between differential *RFX3* motif accessibility and differential *RFX3* expression in excitatory neurons.** Points colored by nominal significance in accessibility and/or expression. The x-axis is the $log_2FC$ from differential motif accessibility analysis, and the y-axis is the $log_2FC$ for differential gene expression analysis. Left: ASD comparison. Right: NDD/ASD comparison.

### Supplementary Fig. 12

a

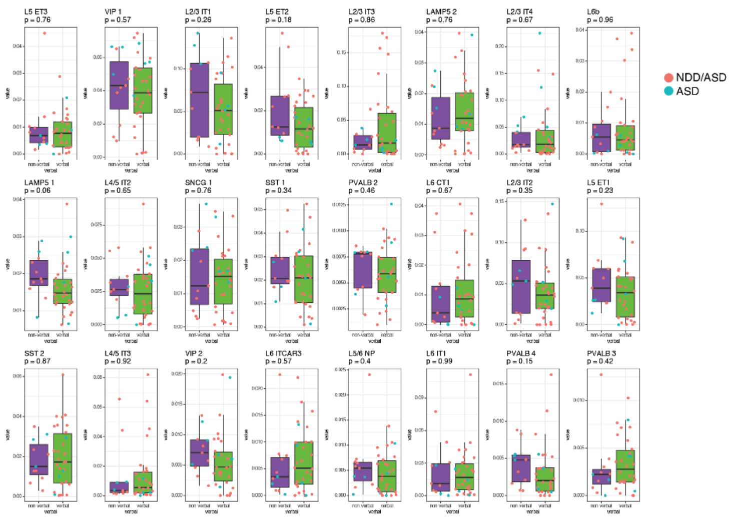

b

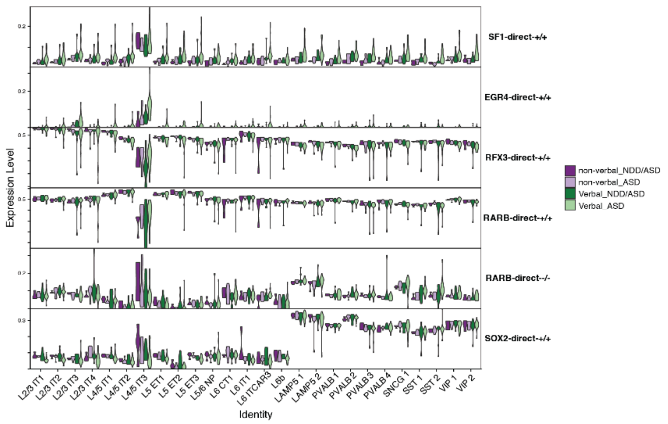

c

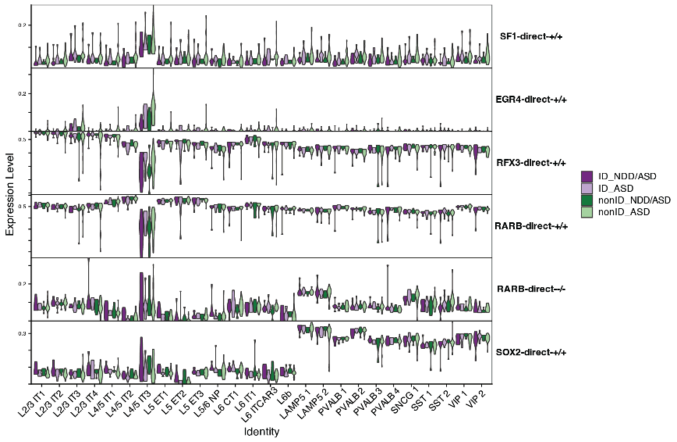

**Supplementary Fig. 12. Cell-type composition and eRegulon regulatory activity across ASD verbal phenotypes.** **a**, Relative proportions per donor of all neuronal clusters in non-verbal versus verbal ASD cohorts. Patients from NDD/ASD are shown in red dots, and ASD patients are shown in blue dots. P-values were calculated using two-sided Mann-Whitney U tests. **b**, Cell-type–specific eRegulon activity distributions for selected transcription factors (*SF1*, *EGR4*, *RFX3*, *RARB*, and *SOX2*) stratified by verbal and ASD phenotype. **c**, Cell-type–specific eRegulon activity distributions for selected transcription factors (*SF1*, *EGR4*, *RFX3*, *RARB*, and *SOX2*) stratified by ID and ASD phenotype.

### Supplementary Fig. 13

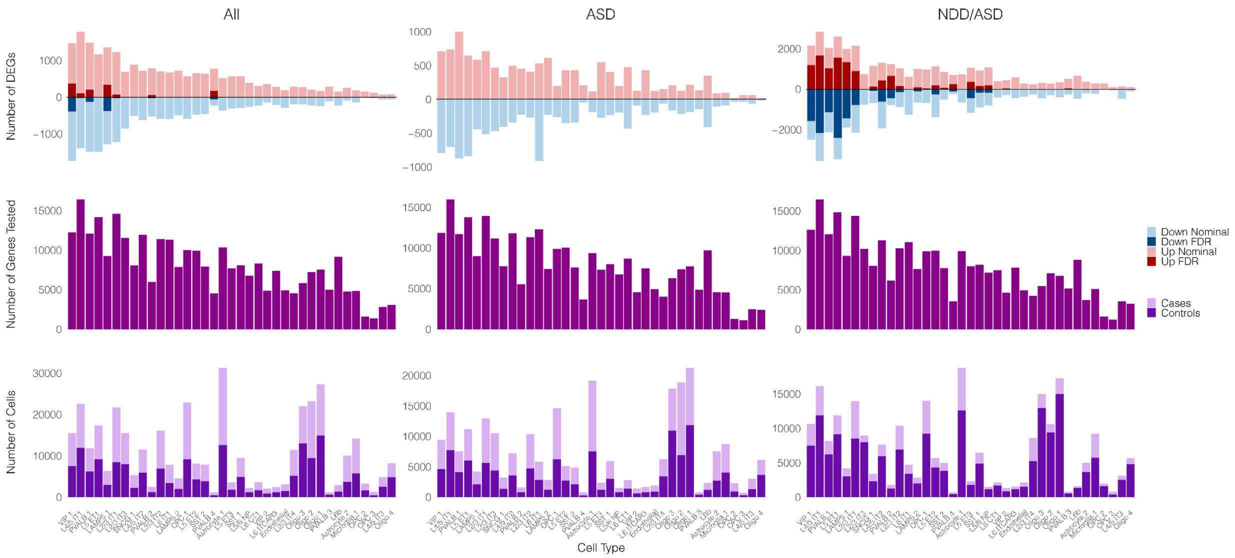

**Supplementary Fig. 13. Differentially expressed genes and cell representation across cell types.** Bar plots summarizing differential gene expression and cell representation across cell types for three comparisons (“All”, “ASD”, and “NDD/ASD”). The top row shows the number of differentially expressed genes (DEGs) per cell type, with upregulated genes (positive values; red shades) and downregulated genes (negative values; blue shades), separated into FDR-significant (darker shades) and nominally significant (lighter shades). The middle row shows the total number of genes tested per cell type. The bottom row shows the number of cells per cell type, split by cases (light purple) and controls (dark purple). Cell types are shown along the x-axis.

### Supplementary Fig. 14

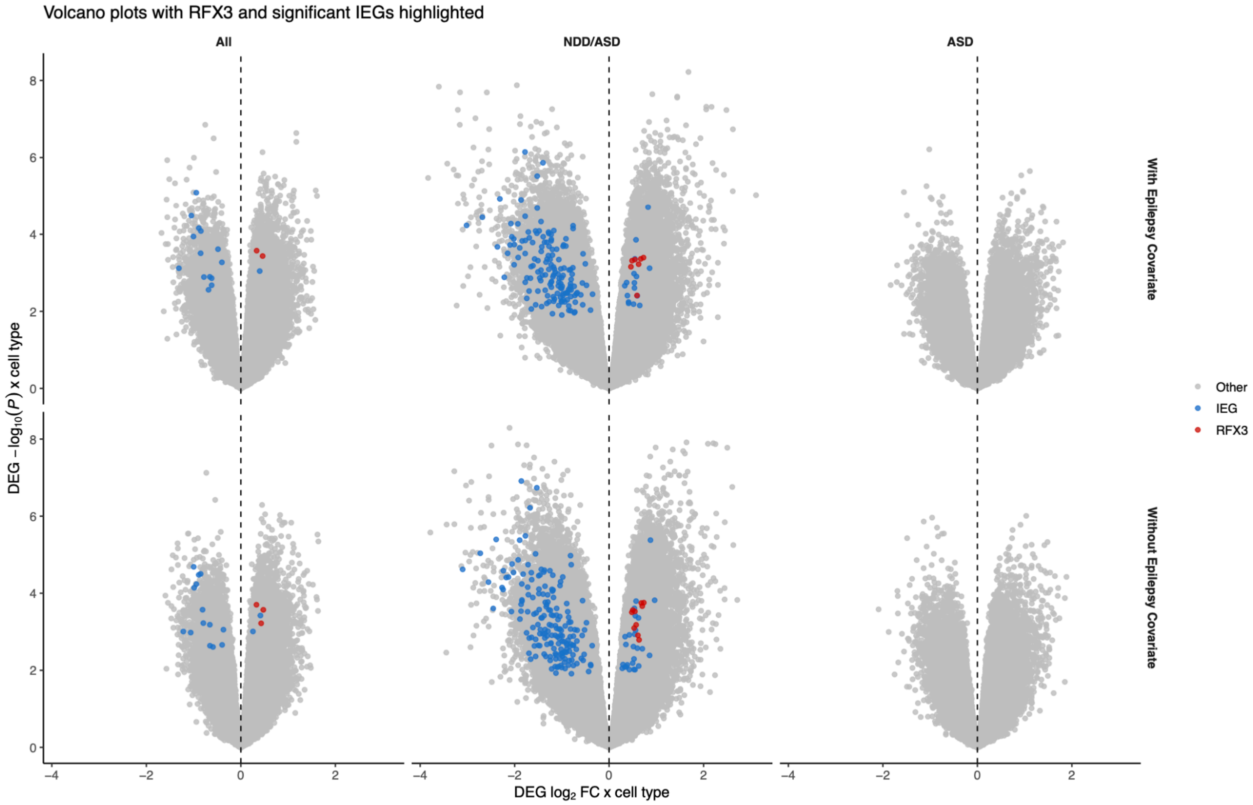

**Supplementary Fig. 14. Volcano plots of differential gene expression highlighting *RFX3* and significant immediate early genes (IEGs).** Six volcano plots show differential expression results across conditions and analysis models. Columns represent All cases and controls, NDD/ASD, and ASD-only subsets. Rows indicate models with (top) and without (bottom) epilepsy added as a covariate. Each point corresponds to a gene by cell type; the x-axis shows $\log_2$ fold change ($\log_2$FC) and the y-axis shows $-\log_{10}$(p-value). Differentially expressed genes (FDR < 0.05) classified as immediate early genes (IEGs) are highlighted in blue, and *RFX3* is highlighted in red. All other genes are shown in gray. The dashed vertical line marks no change ($\log_2$FC = 0). Only statistically significant genes are emphasized.

### Supplementary Fig. 15

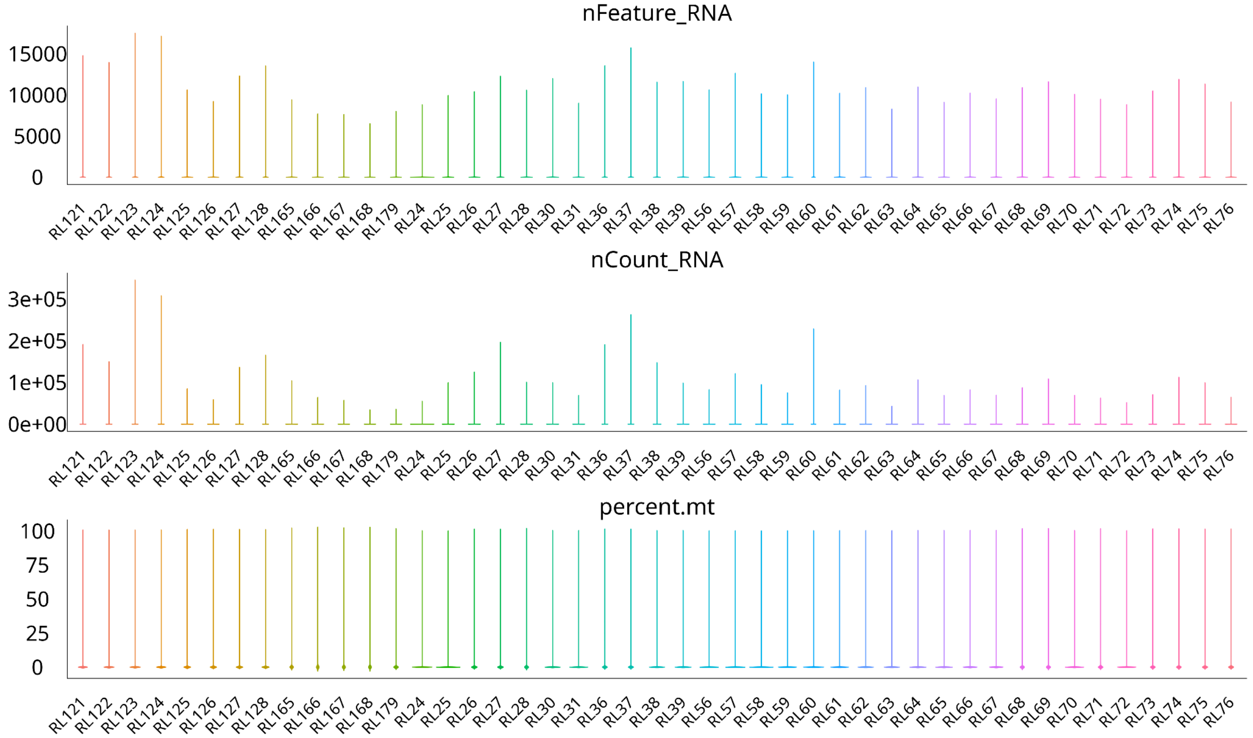

**Supplementary Fig. 15. RNA quality control metrics per well prior to filtering.** Violin plots showing distributions of RNA sequencing quality metrics across wells (RL121–RL76) prior to quality control filtering. Top row: number of detected genes per cell (nFeature_RNA). Middle row: total UMI counts per cell (nCount_RNA). Bottom row: percentage of mitochondrial gene expression (percent_mt). Each violin represents the distribution of values across cells within a given well.

### Supplementary Fig. 16

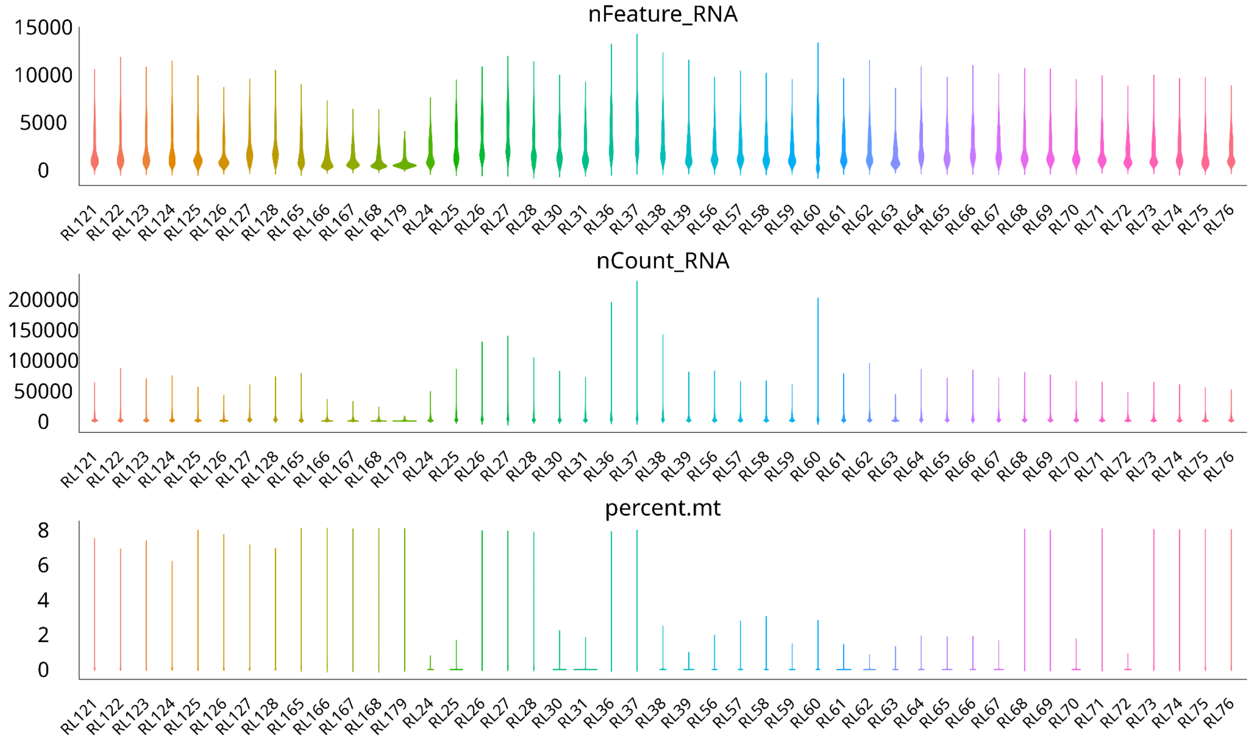

**Supplementary Fig. 16. RNA quality control metrics per well post filtering.** Violin plots showing distributions of RNA sequencing quality metrics across wells (RL121–RL76) after quality control filtering. Top row: number of detected genes per cell (nFeature_RNA). Middle row: total UMI counts per cell (nCount_RNA). Bottom row: percentage of mitochondrial gene expression (percent_mt). Each violin represents the distribution of values across cells within a given well.

### Supplementary Fig. 17

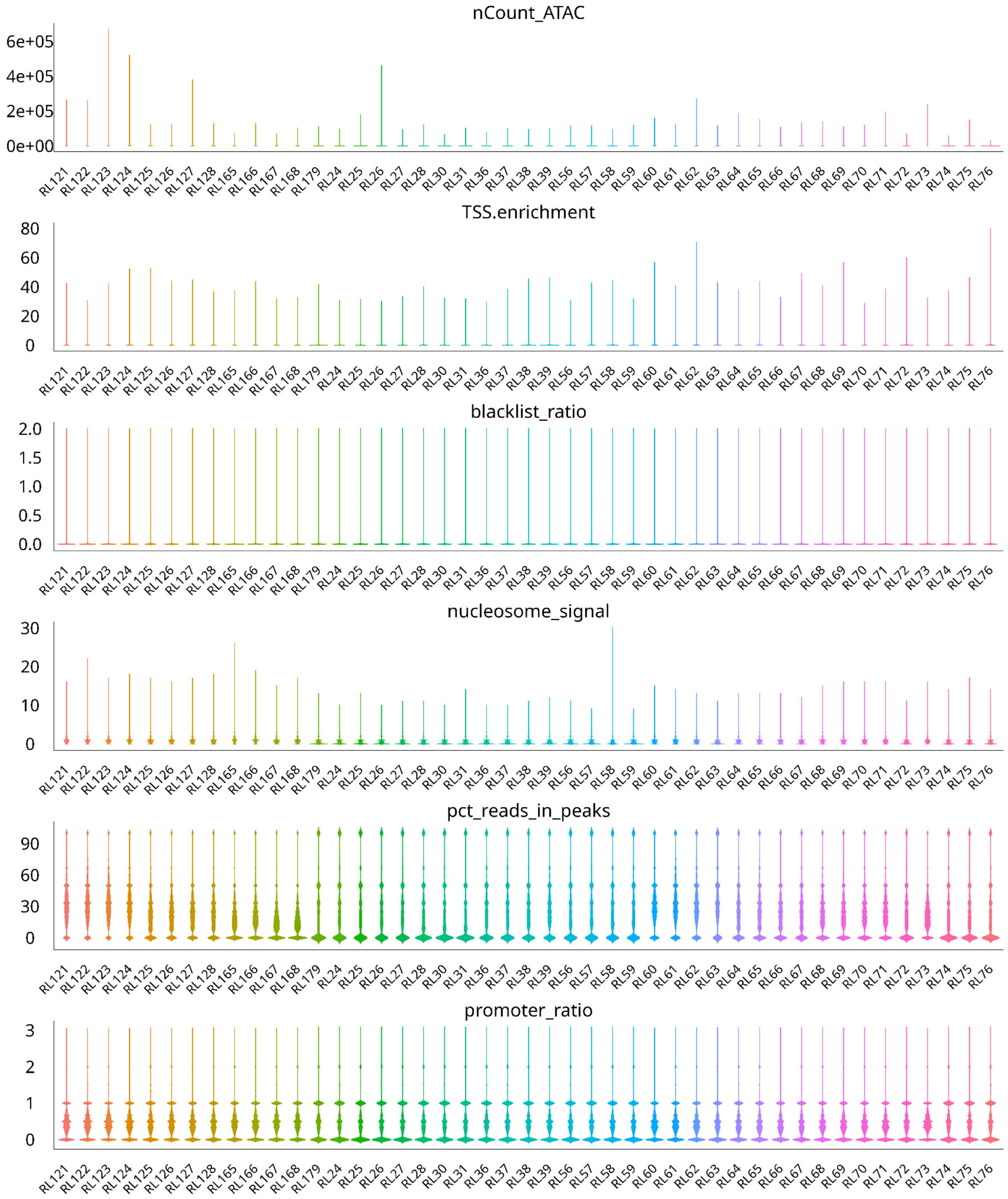

**Supplementary Fig. 17. ATAC-seq quality control metrics per well prior to filtering.** Violin plots showing distributions of ATAC-seq quality metrics across wells (RL121–RL76) prior to quality control filtering. Row display (top to bottom): total ATAC fragment counts per cell (nCount_ATAC), transcription start site enrichment (TSS enrichment), fraction of reads in blacklist regions (blacklist_ratio), nucleosome signal, fraction of reads in peaks (pct_reads_in_peaks), and promoter accessibility ratio (promoter_ratio). Each violin represents the distribution of values across cells within a given well.

### Supplementary Fig. 18

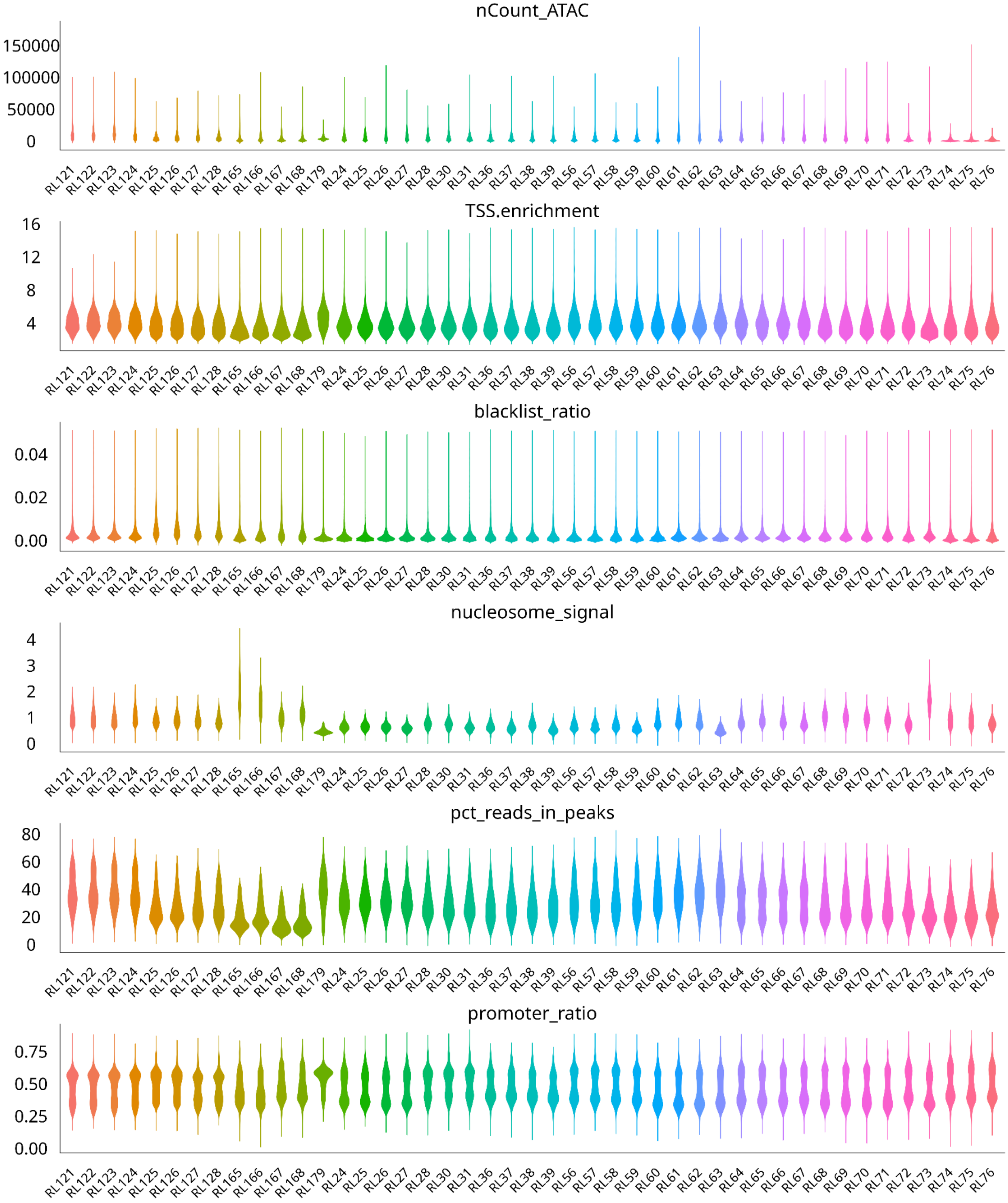

**Supplementary Fig. 18. ATAC-seq quality control metrics per well post filtering.** Violin plots showing distributions of ATAC-seq quality metrics across wells (RL121–RL76) after quality control filtering. Row display (top to bottom): total ATAC fragment counts per cell (nCount_ATAC), transcription start site enrichment (TSS enrichment), fraction of reads in blacklist regions (blacklist_ratio), nucleosome signal, fraction of reads in peaks (pct_reads_in_peaks), and promoter accessibility ratio (promoter_ratio). Each violin represents the distribution of values across cells within a given well.

### Supplementary Fig. 19

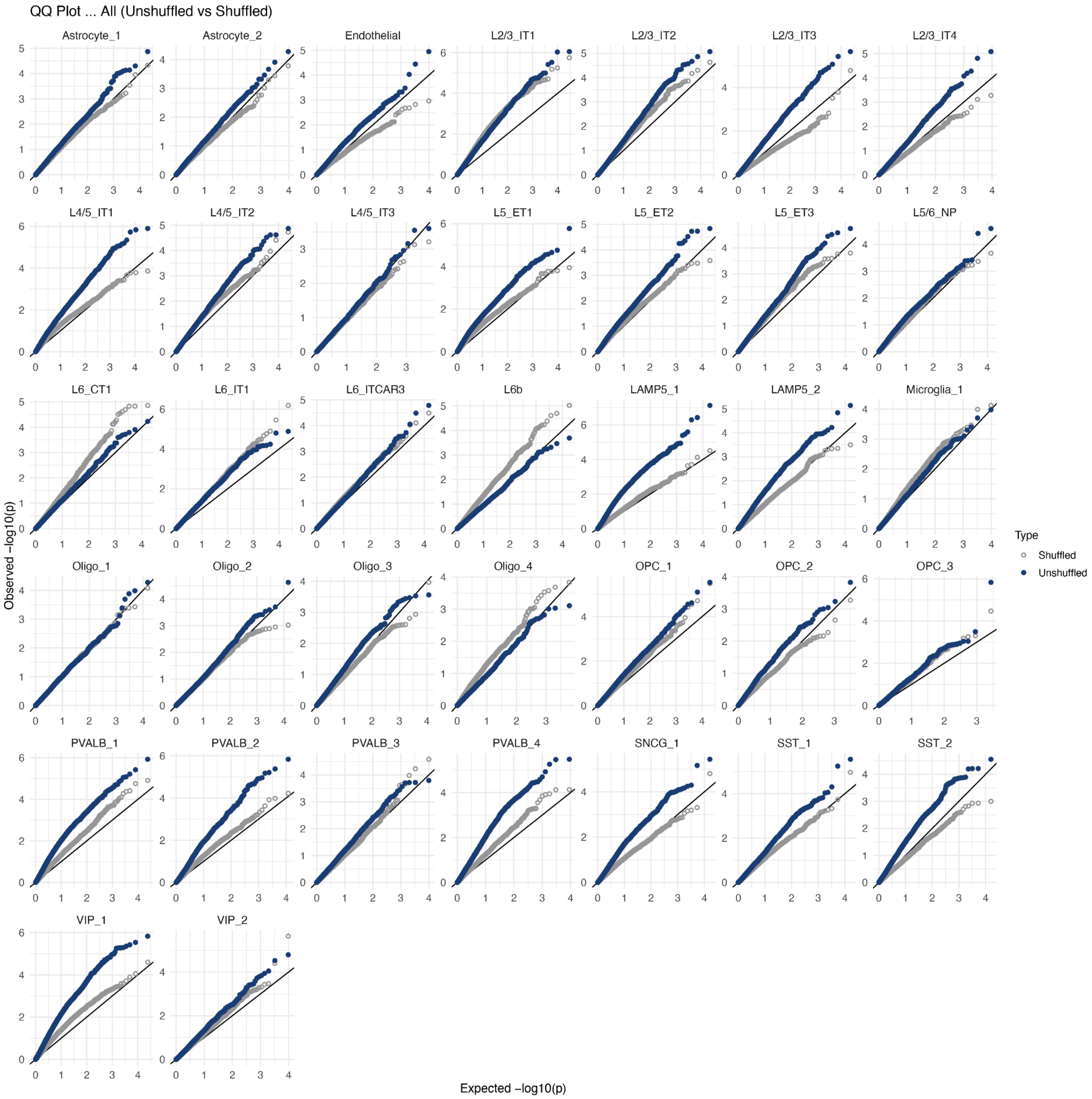

**Supplementary Fig. 19. QQ plots of differential expression p-values across cell types.** Quantile-quantile (QQ) plots comparing observed versus expected $-\log_{10}$(p-values) for differential expression tests across cell types in the “All” comparison. Each panel corresponds to a cell type or subtype. Blue points show results from the true (unshuffled) case-control labels, while gray points show results after random shuffling of case/control labels (null distribution). The diagonal line indicates the expected distribution under the null hypothesis.

### Supplementary Fig. 20

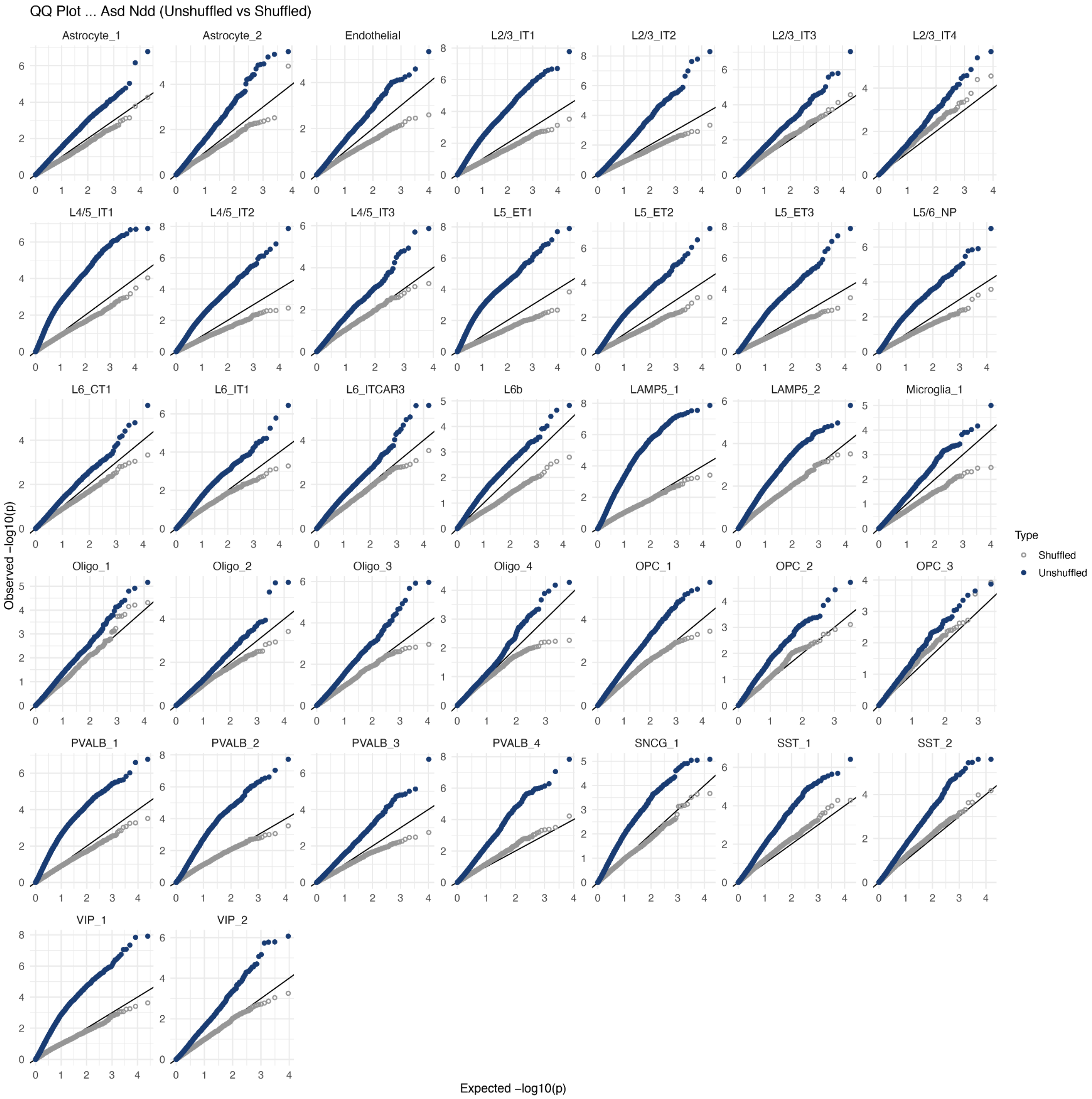

**Supplementary Fig. 20. QQ plots of differential expression p-values across cell types.** Quantile-quantile (QQ) plots comparing observed versus expected $-\log_{10}$(p-values) for differential expression tests across cell types in the “NDD/ASD” comparison. Each panel corresponds to a cell type or subtype. Blue points show results from the true (unshuffled) case-control labels, while gray points show results after random shuffling of case/control labels (null distribution). The diagonal line indicates the expected distribution under the null hypothesis.

### Supplementary Fig. 21

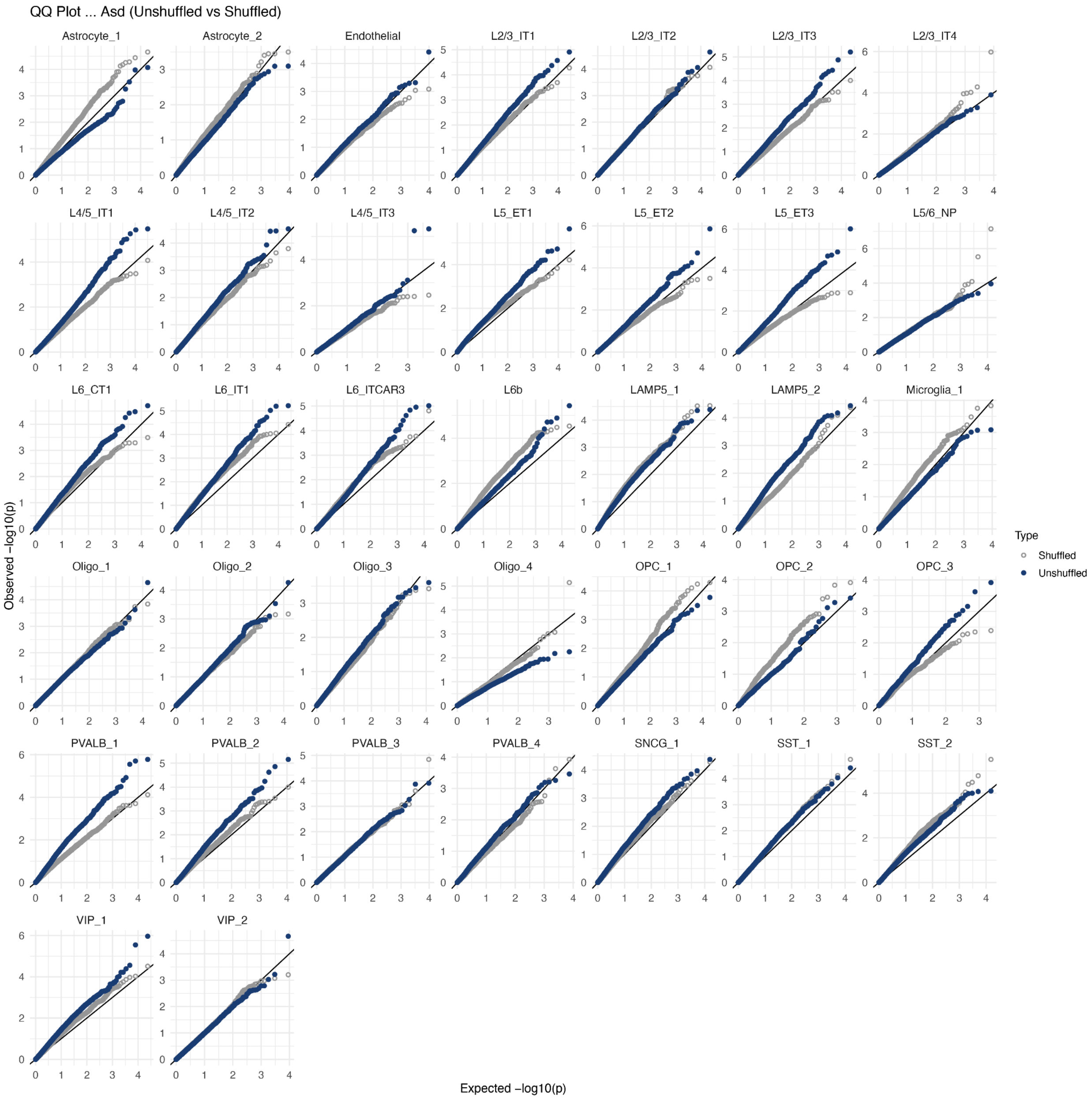

**Supplementary Fig. 21. QQ plots of differential expression p-values across cell types.** Quantile–quantile (QQ) plots comparing observed versus expected $-\log_{10}$(p-values) for differential expression tests across cell types in the "ASD" comparison. Each panel corresponds to a cell type or subtype. Blue points show results from the true (unshuffled) case-control labels, while gray points show results after random shuffling of case/control labels (null distribution). The diagonal line indicates the expected distribution under the null hypothesis.
